# Rbfox1 and *mir-33* regulate pleiotropic roles of JAK/STAT signalling during adult myogenesis in *Drosophila melanogaster*

**DOI:** 10.1101/2024.12.02.626327

**Authors:** Amartya Mukherjee, Upendra Nongthomba

## Abstract

The JAK/STAT pathway is a conserved metazoan signal transduction system with roles in regulating growth, stem cells, and immune responses. Although its components are ubiquitously expressed during development, JAK/STAT signalling exerts cell-specific effects through mechanisms largely unknown. Here, we demonstrate that cell-specific effects of JAK/STAT signalling could be realised through post-transcriptional regulation. We show that JAK/STAT signalling in early adult myogenesis is critical for the patterning of the dorsal longitudinal muscles in *Drosophila melanogaster*. RNA-binding Fox protein 1 (Rbfox1) and the microRNA *mir-33* regulate the transcript stability of *Signal-transducer and activator of transcription protein at 92E* (*Stat92E*), which encodes the sole *Drosophila* STAT orthologue. Moreover, Rbfox1 also controls the alternative splicing of *Stat92E* and its key downstream effector, *Zn finger homeodomain 1* (*zfh1*). The coupling of Rbfox1 with Stat92E function, during early adult myogenesis, not only allows maintenance of stemness, but also mediates filamentous actin dynamics, and prevents apoptosis in myoblasts via some other Stat92E target genes. Given that the functions of the JAK/STAT pathway in cell proliferation and survival are conserved between *Drosophila* and vertebrates, our report presents a novel example for the context-dependant regulation of the developmental role of an important signalling pathway.

## Introduction

The JAK/STAT pathway is a conserved signalling system found in all metazoans. It is activated by a cytokine binding to its receptor on the cell surface, triggering a cascade of events that leads to the transcription of target genes (Herrera and Bach 2019). Although signalling pathways, transmitting information between cells, coordinating their interactions, and leading to organogenesis, act recurrently at different times, and in different regions of the embryo (Sanz-Ezquerro et al. 2017), how cells modulate and respond to the JAK/STAT pathway in cell type-specific ways remains a puzzle. The fruit fly *Drosophila melanogaster* has a complete, but relatively simple, JAK/STAT system, consisting of three ligands, Unpaired 1 (Upd1), Upd2, and Upd3; one receptor, Domeless (Dome); one Jak family tyrosine kinase (JAK), Hopscotch (Hop); and one STAT protein, Signal-transducer and activator of transcription protein at 92E (Stat92E) (Herrera and Bach 2019). This makes *D. melanogaster* an excellent model for developmental genetic investigations of the many roles of the JAK/STAT pathway, with important ramifications for vertebrate model organisms.

An ideal model system to investigate mechanisms of morphogenesis are the large indirect flight muscles (IFMs) that span the entire *Drosophila* thorax (Gunage et al. 2017; Rai et al. 2014). Like all adult fly muscles, the IFMs are formed from a pool of undifferentiated myoblasts, called adult muscle precursors, during metamorphosis (Currie and Bate 1991). *D. melanogaster* IFMs comprise, per hemithorax, seven dorsoventral flight muscles (DVMs), and six dorsal longitudinal flight muscles (DLMs). While the DVMs form *de novo* from fusion of founder cells (FCs) and fusion-competent myoblasts (FCMs), the DLMs are formed through fusion of waves of FCMs to three persistent larval muscles templates that escape histolysis, the larval oblique muscles (LOMs), that function as FCs, making DLMs similar to vertebrate skeletal muscles in their development and organisation (Dutta et al. 2004; Fernandes et al. 1991; Taylor 2013). Because the final number of DLM fascicles (muscle fibre bundles) is determined by the splitting of the templates, which is directly controlled by the fusion of myoblasts (Fernandes and Keshishian 1996), any disruptions to the specification, proliferation, migration, and fusion of myoblasts can severely affect DLM biogenesis.

In this study, we report that the cascade of JAK/STAT pathway signalling events is responsible for muscle formation. Moreover, we show that differential expression of RNA-binding Fox protein 1 (Rbfox1) and the microRNA *mir-33* during DLM formation regulates the temporal expression dynamics of Stat92E and its target Zn finger homeodomain 1 (Zfh1), a transcriptional repressor which promotes stem cell self-renewal (Herrera and Bach 2019). Rbfox1, in its role as a splicing regulator, directly dictates the conserved switch between the splice variants of Stat92E and Zfh1, highlighting the robustness of the regulatory mechanisms. Thus, downregulation of Rbfox1 and components of the JAK/STAT signalling pathway perturbs splitting of the larval muscle templates. Together, our findings reveal that distinct facets of Rbfox1 function, during early adult myogenesis in *Drosophila*, govern tissue-specific aspects of JAK/STAT signalling, including maintenance of stemness, mediation of filamentous actin (F-actin) dynamics, and inhibition of apoptosis in myoblasts.

## Results

### Rbfox1 function is required in myogenic cells only for normal patterning of the DLMs

RNA-binding Fox protein 1 (Rbfox1), previously known as Ataxin 2-binding protein 1 (A2bp1), has myriad functions: it is a transcriptional co-activator (Shukla et al. 2017), a tissue-specific splicing factor (Underwood et al. 2005), and affects mRNA stability and translation (Carreira-Rosario et al. 2016). Recent research in our laboratory has elucidated the role of *Drosophila* Rbfox1 in muscle development (A Mukherjee and U Nongthomba, under revision; Nikonova et al. 2022). We have shown that it is crucial for specifying muscle fibre fate, and for maintaining the stoichiometry of sarcomeric proteins (Nikonova et al. 2022).

Based on these key findings, we decided to shift our attention to its role in early stages of adult muscle development (myoblast proliferation and fusion). First, we decided to discern the cell-autonomous and/or non-autonomous requirements for *Rbfox1* in this process. The appropriate formation of the six DLM fascicles is dictated by spatial and temporal cues emanating from the myoblasts, the templates, the epidermis, and the metamorphosing innervation (Roy and VijayRaghavan 1998). To this end, when we knocked-down *Rbfox1* using the pan-neuronal driver *elav-GAL4* (Lin and Goodman 1994), due to the strong induction of RNAi, we observed embryonic lethality, which agrees with its important role in the development of the nervous system (reviewed in Mukherjee and Nongthomba 2023). On the other hand, pan-neuronal *Rbfox1* knock-down using *nSyb-GAL4* (Pauli et al. 2008) produced viable flies, but they had completely compromised flight ability (Fig. 1A). Likewise, knock-down of *Rbfox1* with *sr-GAL4* (Ghazi et al. 2000), which drives expression in muscle attachment sites, resulted in a major loss of flight ability as compared to controls (Fig. 1A). However, when the hemithoraces were observed under polarised light, despite *Rbfox1* knock-down, adults showed presence of typical six DLMs, similar to those of controls (Fig. 1B). These data suggest that *Rbfox1* function in neurons and tendon cells is essential to confer normal flight ability, but is dispensable for DLM template splitting.

**Figure 1.**
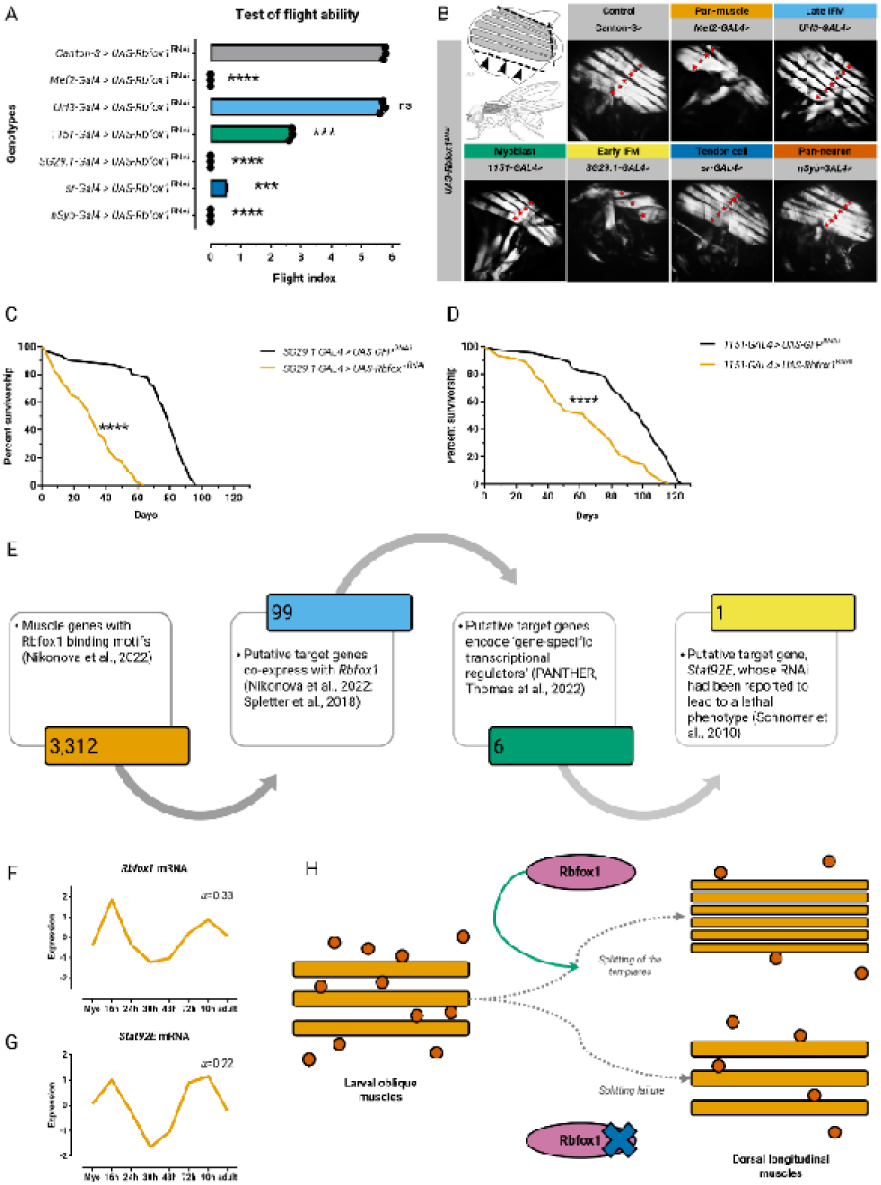
*Rbfox1* expression during early stages of myogenesis is necessary for splitting of the DLM templates. **(A)** Quantification of flight ability after *Rbfox1* knockdown. Genotypes as noted. Significance is from paired *t* test (\*\*\**P* < 0.001; \*\*\*\**P* < 0.0001). **(B)** Polarised microscopy images of hemithorax from flies. Genotypes as noted. Red stars indicate DLM fascicles. **(C, D)** Survival curves at 22°C. Genotypes as noted. Significance is from Log-rank test (\*\*\*\**P* < 0.0001). **(E)** Bioinformatic work-flow leading to the identification of *Stat92E* as a putative target of *Rbfox1.* Standard normal count values for *Rbfox1* **(F)** and *Stat92E* **(G)** from an mRNA-seq developmental time-course of wild-type IFMs (Spletter et al., 2018). **(H)** Graphical summary.

The constitutive knock-down of *Rbfox1* with *Mef2-GAL4* (Ranganayakulu et al. 1998) resulted in embryonic or larval lethality, which may have been influenced by defects in larval muscles. Thus, to directly address its role during adult myogenesis, and to reduce *Rbfox1* levels in myoblasts at early adult myogenesis stage, right when DLM development is initiated, we used a α*Tub84B-GAL80^ts^* transgene (*tub-GAL80^ts^*, McGuire et al. 2003) in addition to the *Mef2-GAL4* transgene. *tub-GAL80^ts^;Mef2-GAL4>UAS-Rbfox1^RNAi^*progeny were raised at the permissive temperature of 18°C till late third instar larval stage, and shifted to the restrictive temperature thereafter. Progeny flies were found to be entirely incapable of flight (Fig. 1A). More strikingly, muscle birefringence revealed that they had severe muscle defects: approximately 70% of such flies had DLMs with template splitting defects, compared to the hemithoraces of controls which showed the presence of the typically six DLM fascicles (Fig. 1B). Also, whereas roughly 80% of the flies with *Rbfox1* knock-down had a hypercontraction muscle phenotype, in which the fascicles appeared to be broken and pulled toward the attachment sites, no hypercontraction was seen in controls (Nikonova et al. 2022; Nongthomba et al. 2003). The observation of two distinct muscle phenotypes, splitting defects and hypercontraction, suggests that *Rbfox1* is involved in two independent processes: template splitting and myofibrillogenesis. This is consistent with the transcriptomics resource of developing flight muscles (Spletter et al. 2018), according to which the *Rbfox1* mRNA exhibits a bimodal temporal expression profile: the mRNA level peaks in myoblasts between 16 h to 24 h APF, is significantly downregulated between 30 h to 72 h APF, and strongly increases until 90 h APF.

### Rbfox1 *expression during early stages of myogenesis is necessary for splitting of the DLM templates*

Reasoning that *Rbfox1* contributes to patterning of the DLMs at early myogenesis stages, we performed ‘early’ knock-down of *Rbfox1* driven by *1151-GAL4* (Roy and VijayRaghavan 1997) and *SG29.1-GAL4* (Roy et al. 1997), and found that progeny flies had significantly lower flight ability compared to the control (Fig. 1A), and abnormal numbers (three or four) of DLM fascicles per hemithorax (Fig. 1B). Specifically, knock-down of *Rbfox1* driven by *SG29.1-GAL4* elicits a severe hypercontraction phenotype, similar to that produced by *Mef2-GAL4* (Fig. 1B), uncovering the later role of Rbfox1 in myofibrillogenesis (Nikonova et al. 2022). These data validated our hypothesis that indeed *Rbfox1* function during DLM splitting lies within the temporal window of early myogenesis (Fig. 1F).

To validate that the expression of *Rbfox1* in early IFM development is sufficient to dictate the generation of the proper DLM pattern, we performed ‘late’ knock-down of *Rbfox1* using *UH3-GAL4* (Singh et al. 2014). Progeny flies exhibited normal flight ability (Fig. 1A), and had the typical six well-organised DLM fascicles per hemithorax (Fig. 1B). These results suggest that the temporal focus of Rbfox1 function in DLM template splitting lies in early, but not late (myofibre growth), myogenesis stages.

Next, we asked whether the reduction of the number of DLM fascicles, reminiscent of skeletal muscle dysplasia, affects the survival of flies. Indeed, flies with ‘early’ knock-down of *Rbfox1* showed significantly reduced median survival compared to controls (Fig. 1C,D). This finding is consistent with the reduced life expectancy which characterises congenital muscular dystrophies (Zambon and Muntoni 2021).

### Bioinformatic identification of early myogenesis genes with Rbfox1 binding motifs

To gain insight into the underlying cause of the patterning phenotype, we sought to identify Rbfox1 target genes in muscle. In our previous publication, we have reported the results of a bioinformatic search to identify 3,312 genes with Rbfox1 binding motifs (Nikonova et al. 2022). Second, we superimposed the putative Rbfox1 targets against genes expressed in *Drosophila* flight muscles (Spletter et al. 2018), their intersection representing the muscle-expressed genes which may be regulated by Rbfox1 (Supplemental Fig. S1A). Interestingly, we observed that the highest percentage of putative muscle-expressed Rbfox1 target genes (nearly 50%) are found in Mfuzz Clusters 35 and 36, distinct gene expression profile clusters generated from the time point-specific transcriptomes of developing flight muscles, both of which exhibit bimodal temporal expression profiles reminiscent of that of *Rbfox1* itself (Fig. 1D; Supplemental Fig. S1B). Therefore, third, we focused on the genes from the common subset obtained for Cluster 36, which contains *Rbfox1*, reasoning that these 99 genes, with temporal expression dynamics similar to that of *Rbfox1*, may be activated by Rbfox1 itself. Fourth, enrichment analysis was performed with the PANTHER Classification System (Thomas et al. 2022) using available Gene Ontology (GO) terms for *D. melanogaster*. Genes with Rbfox1 motif instances are enriched GO terms related to transcription, muscle development and cytoskeletal organisation, for example, ‘transcription regulator activity’, ‘developmental process’, ‘locomotion’, ‘cell adhesion molecule’, and ‘cytoskeletal protein’, strongly suggesting that genes important for early muscle developmental processes are likely targets of Rbfox1 regulation. We next proceeded to experimentally validate some of these candidate genes.

### Stat92E *is a Rbfox1 target gene during DLM patterning*

Since the observed morphological transition suggests precise temporal transcriptional regulation, we next selected candidate Rbfox1 target genes in Cluster 36 which encode for ‘gene-specific transcriptional regulator’ to verify the direct or indirect involvement of Rbfox1 in the pivotal role transcriptional regulation plays during muscle development. The six mapped PANTHER database hits encode the DNA-binding transcription factors CG42526, and Signal-transducer and activator of transcription protein at 92E (Stat92E); the Cys_2_His_2_ (C2H2) zinc finger transcription factors Zn finger homoeodomain 1 (zfh1), and CG11902; the homeodomain transcription factor Onecut; and the C4 zinc finger nuclear receptor Seven up (Svp) (Fig. 1E; Fig. 5A; Supplemental Fig. S3A). To select the candidate target gene, we filtered the six PANTHER database hits through the results of a *Mef2-GAL4*-driven genome-wide RNAi screen (Schnorrer et al. 2010), and found that *Stat92E* was the only gene whose RNAi had been reported to lead to a lethal phenotype. Notably, *zfh1* is a key direct Stat92E target gene in somatic cyst stem cells (CySCs) (Leatherman and DiNardo 2008). These leads, namely, the presence of *Stat92E*, and its downstream effector, *zfh1*, in the same cluster as *Rbfox1*, and the late pupal lethal phenotype of its RNAi (Schnorrer et al. 2010), merited *Stat92E* being chosen as the promising putative Rbfox1 target gene.

The role of JAK/STAT pathway, in the context of muscle development, was first characterised when it was shown that embryos lacking maternal and zygotic *Stat92E* have completely penetrant defects in somatic muscle development, with many muscles missing, the remaining muscles having no clear identity or shape, and myoblasts remaining unfused (Liu et al. 2009). To establish whether Rbfox1 regulates *Stat92E*, we interrogated whether *Rbfox1* misexpression affects the transcript levels of *Stat92E*. We found that that there is a direct correlation between *Rbfox1* expression and *Stat92E* expression: knock-down of *Rbfox1* results in downregulation of *Stat92E*, whereas overexpression of *Rbfox1* causes upregulation of *Stat92E* (Fig. 2A).

**Figure 2.**
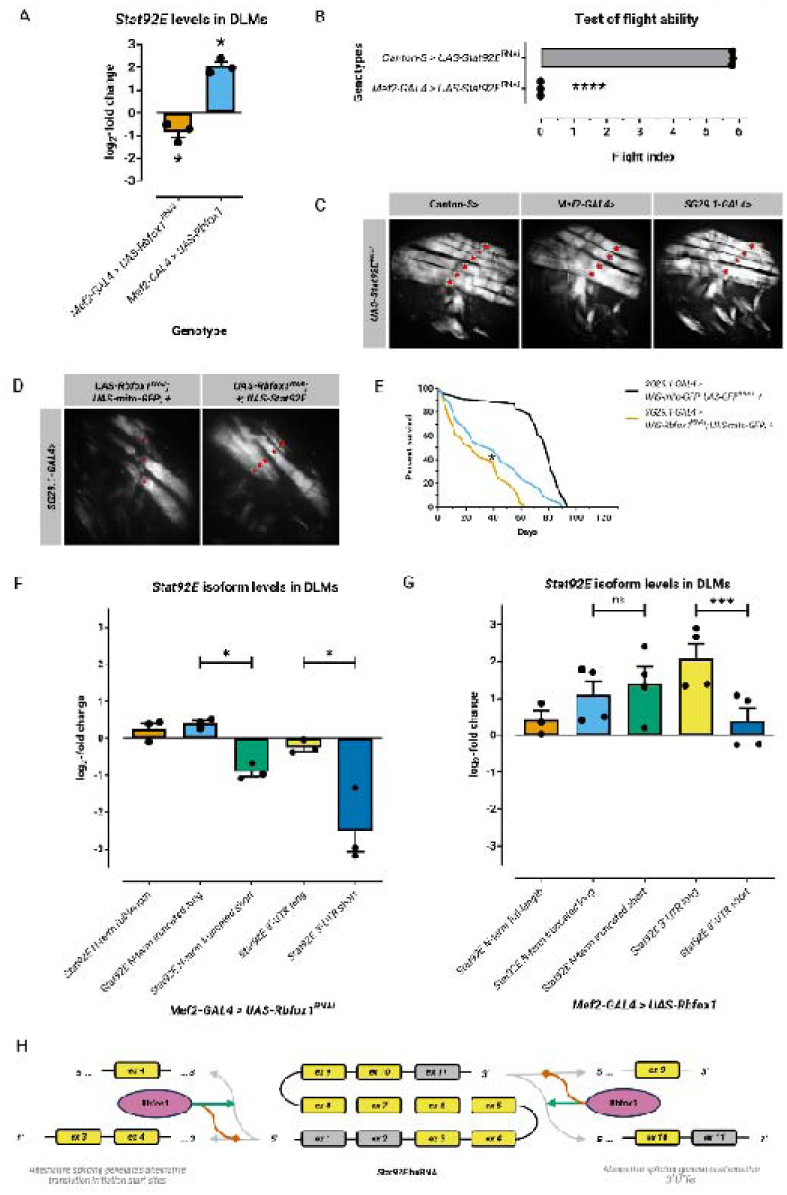
The alternative splicing of Stat92E transcripts by *Rbfox1* is necessary for splitting of the DLM templates. **(A)** Quantification of RT-qPCR data for *Stat92E* transcript levels in IFMs from *Rbfox1* misexpression. Significance is from paired t test (\**P* < 0.05). **(B)** Quantification of flight ability after *Stat92E* knockdown. Genotypes as noted. Significance is from paired *t* test (\*\*\*\**P* < 0.0001). **(C, D)** Polarised microscopy images of hemithorax from flies. Genotypes as noted. Red stars indicate DLM fascicles. **(E)** Survival curves at 22°C. Genotypes as noted. Significance is from Log-rank test (\**P* < 0.05). Quantification of RT-qPCR data for *Stat92E* splice variant levels in IFMs from *Rbfox1* knockdown **(F)** and overexpression **(G)**. Significance is from paired t test (\**P* < 0.05; \*\*\**P* < 0.001). **(H)** Graphical summary.

Next, we determined whether Stat92E is involved in DLM template splitting. While knock-down of *Stat92E* driven by *1151-GAL4*, specific to disc-associated myoblasts, proved to be lethal, that by *Mef2-GAL4* led to viable progeny flies, but they were all flightless (Fig. 2B). Moreover, muscle birefringence revealed that knock-down driven by *Mef2-GAL4* and *SG29.1-GAL4*, produced lesser than normal number of DLM fascicles (Fig. 2C), phenocopying the *Rbfox1* knock-down condition. This suggested that *Stat92E* is required for DLM template splitting, and is possibly regulated by Rbfox1.

To validate this genetic interaction, we ectopically overexpressed *Stat92E* in the genetic background of *Rbfox1* knockdown driven by *SG29.1-GAL4.* We introduced the *UAS-mito-GFP* transgenic construct in the *UAS-Rbfox1^RNAi^* genetic background to control for GAL4 dosage. Muscle birefringence revealed that *Stat92E* is sufficient to rescue the splitting defect cause due to downregulation of *Rbfox1* (Fig. 2D). The *Stat92E* expression also led to significant lifespan extension compared to the *Rbfox1^RNAi^* control (Fig. 2E). Interestingly, however, *Stat92E* failed to prevent the hypercontraction phenotype typical of the *Rbfox1^RNAi^*control (Fig. 2D), suggesting that, during its second expression peak, *Stat92E*, unlike *Rbfox1*, is not involved in myofibrillogenesis.

### *Rbfox1 regulates the alternative splicing of* Stat92E *transcripts*

*Stat92E* is the single *Drosophila* orthologue of the seven mammalian *STAT* genes (Herrera and Bach 2019). Several structural features conserved in STAT proteins can also be recognised in Stat92E, being organised into functional modular domains. The longest isoform, Stat92E-PK, comprises of the amino terminal domain (N-domain), the adjacent coiled-coil domain, a DNA-binding domain in the central region, a linker domain, an Src homology 2 (SH2) domain, and the carboxyl terminal domain (Chen et al. 1998). In addition to the full-length form (STATα) (*e.g.*, Stat92E-PK, length 818 aa, mass 92,573 Da), STAT proteins can also be present as C-terminally truncated forms (*e.g.*, Stat92E-PL, length 679 aa, mass 76,791 Da) generated by alternative splicing (STATβ) or by proteolytic processing (STATγ) (Mitchell and John 2005). Although the truncated STATβ, which lacks the C-terminal activation domain, was earlier considered as a dominant negative form (Caldenhoven et al. 1996), studies have indicated specific functions for the STATβ isoform (Maritano et al. 2004; Xia et al. 2001; Yoo et al. 2002).

It is established that the presence of the Rbfox1 consensus binding site in the region upstream or downstream of the alternative exon is an indication that they are likely to be directly regulated by Rbfox1 (Pedrotti et al. 2015). In a dataset of neuronal genes involved in cell differentiation and proliferation, RBFOX1 was reported to change the alternative splicing events involving *STAT3* (Fogel et al. 2012). Interestingly, there are two highly conserved Rbfox1 binding sites (core *GCAUG* sites, with the variable 5’ *U* residue) on the intron preceding the alternative exons at the 3’ end, and a *UGCAUG* motif on the intron between the alternative exons at the 5’ end of *Stat92E* (Supplemental Fig. S3F). The alternative splicing of *STAT* paralogues is conserved. Specifically, while at least one Rbfox1 binding site at the *STAT* 3’ end is conserved from *Caenorhabditis elegans* to human, the 5’ end binding site is absent in *C. elegans*, but is conserved from *D. melanogaster* to human (Supplemental Fig. S3C,D), suggesting that the 5’ end alternative splicing is a more recent modification of the *Stat92E* transcript than the 3’ end one.

To test the hypothesis that Rbfox1 is involved in the alternative splicing of the *Stat92E* transcript, we determined the expression levels of the various *Stat92E* splice variants under conditions of *Rbfox1* misexpression. In case of the 3’ end alternative splicing, we found that *Rbfox1* knock-down causes a selective downregulation of the *3’-UTR-short* variant, whereas *Rbfox1* overexpression causes the preferential upregulation of the *3’-UTR-long* variant (Fig. 2F). This suggests that Rbfox1 facilitates the switch from the default *3’-UTR-short* to the *3’-UTR-long* variant (Fig. 2H). In case of the 5’ end alternative splicing, we found that *Rbfox1* knock-down causes a selective downregulation of the *5’-UTR-truncated short* variant, whereas *Rbfox1* overexpression causes its preferential upregulation, compared to the *5’-UTR-full* and *5’-UTR-truncated long* variants (Fig. 2G,H). Together, these findings suggest that the indeed the splice variant-specific expression of *Stat92E* is regulated by Rbfox1, with the switch being critical for DLM development and function.

We predicted the structural differences between the Stat92E isoforms encoded by the splice variants under native conditions. Although there are no notable differences between the short (Stat92E-PK) and long (*e.g*., Stat92E-PL) isoforms at their C-terminal region, there are stark differences between their N-terminal regions: the short isoform lacks the N-terminal protein-protein interaction domain (Pfam entry: PF02865) (Supplemental Fig. S4A,B), which is known to strengthen interactions between STAT dimers on adjacent DNA-binding sites. Furthermore, the short isoform has a higher score log-likelihood ratio (LLR), suggesting that it is more disordered than the long isoform. While the long isoform is structured between its first hundred residues (indicated by a peak in the-4*PAPA), representing the STAT dimerisation domain, the short isoform abruptly commences with a unstructured, prion-like domain (PrLD), representing the, perhaps truncated, coiled-coil STAT domain, which is implicated in protein-protein interactions (Supplemental Fig. S4C,D). Based on these data, we propose that the short Stat92E isoform, lacking the dimerisation domain, which promotes cooperativity of STAT binding to its DNA binding sites, may behave in a dominant negative fashion.

### Rbfox1 regulates *Stat92E* transcript stability by preventing *mir-33* binding

The 1.438 kb-long 3’-UTR of the *Stat92E 3’-UTR-long* variant, but not the 0.393 kb-long *3’-UTR-short* one, bears a few conserved Rbfox1 binding sites (Supplemental Fig. S3C,F). Interestingly, the ***UGCA****UG* Rbfox1 binding site overlaps with a conserved *mir-33-5p* site, *CAA**UGCA*** (Supplemental Fig. S3F). Moreover, *mir-33* binding sites are also found on the *Rbfox1* transcripts with longer 3’-UTR (Supplemental Fig. S3E). Lee and coworkers (2016) have reported that Rbfox1 and microRNA (miRNA) binding sites overlap significantly, and Rbfox1 binds to the 3’-UTR of target mRNAs in the cytoplasm of neurons, increasing their stability and translation, and antagonising miRNA-mediated degradation.

To identify the role of *miR-33* in adult myogenesis, we perturbed the expression of *miR-33* driven by *tub-GAL80^ts^;Mef2-GAL4* and *SG29.1-GAL4.* Overexpression of *miR-33* resulted in pupal lethality, and adversely affected flight ability, whereas its depletion had no such effects (Fig. 3A,B,D). Similarly, while the hemithoraces of flies with *mir-33* depletion showed presence of typical six DLMs, those with *mir-33* overexpression had an abnormal number of four DLM fascicles (Fig. 3C,E). These data suggest that DLM template splitting is robust to downregulation of *mir-33* but is hindered by its expression exceeding a certain threshold.

**Figure 3.**
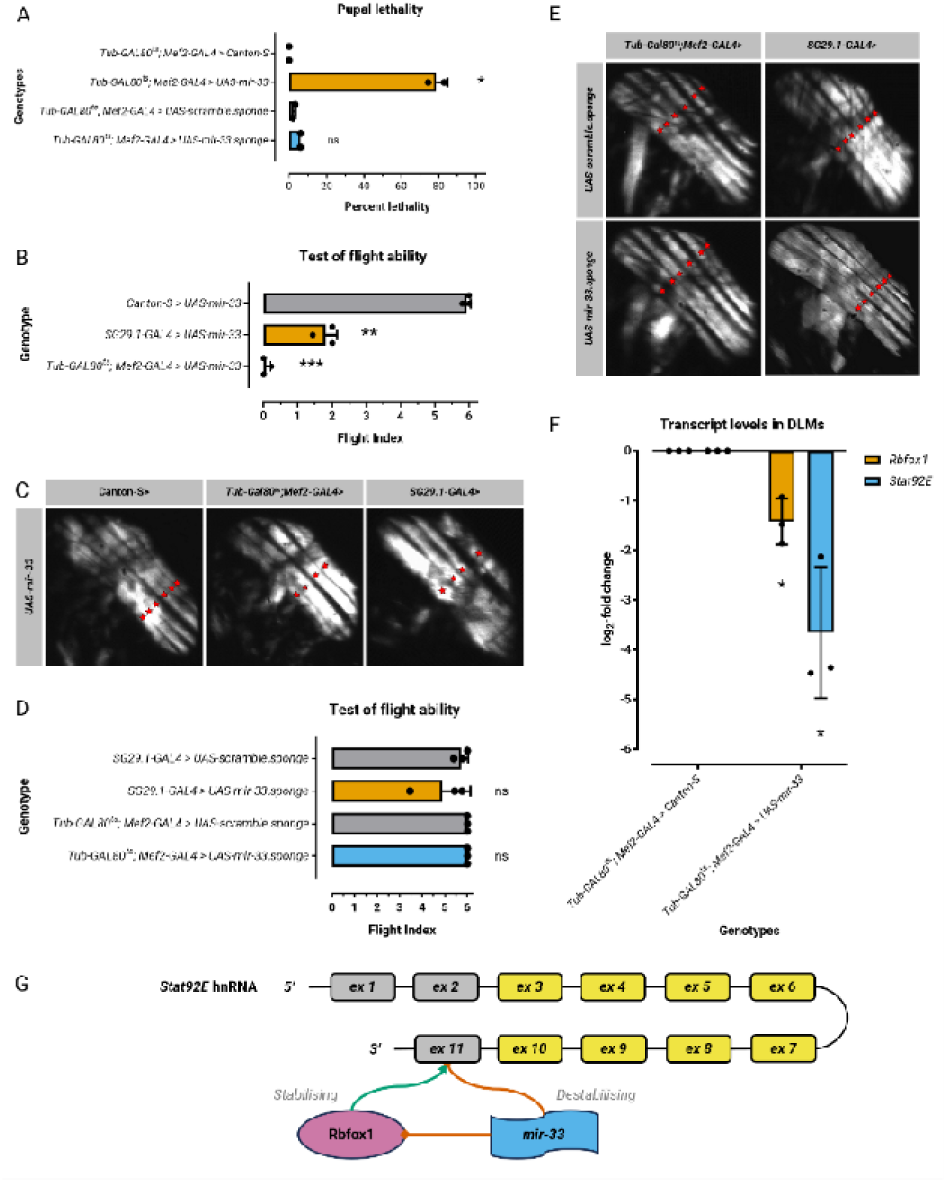
Rbfox1 regulates *Stat92E* transcript stability by preventing *mir-33* binding. **(A)** Quantification of pupal lethality from *mir-33* misexpression. Genotypes as noted. Significance is from paired t test (\**P* < 0.05). Quantification of flight ability after *mir-33* depletion **(B)** and overexpression **(D)**. Genotypes as noted. Significance is from paired *t* test (\*\**P* < 0.01; \*\*\**P* < 0.001). **(C, E)** Polarised microscopy images of hemithorax from flies. Genotypes as noted. Red stars indicate DLM fascicles. **(F)** Quantification of RT-qPCR data for *Rbfox1* and *Stat92E* transcript levels in IFMs from *mir-33* overexpression. Significance is from paired t test (\**P* < 0.05). **(G)** Graphical summary.

To establish whether *mir-33* regulates *Rbfox1* and *Stat92E*, we interrogated whether its overexpression affects their transcript levels. We found that that *mir-33* expression indeed results in downregulation of *Rbfox1* and, more strongly, *Stat92E* (Fig. 3F). Hence, we propose that during adult myogenesis, Rbfox1 occupies the 3’-UTR of *Stat92E* transcripts, thereby preventing the binding of *mir-33* to their shared binding site (Fig. 3G). Thus, naturally, the Rbfox1-*Stat92E* protein-RNA interaction remains impervious to reduction in *mir-33* levels. However, a drastic increase in *mir-33* levels not only reduces *Rbfox1* expression but outcompetes Rbfox1 for binding to the *Stat92E* 3’-UTR as well as, thereby doubly affecting *Stat92E* expression.

To support our proposition using bioinformatic analysis, we superimposed the putative *mir-33* targets (TargetScanFly, Agarwal et al. 2018) against the distinct expression profile clusters (Spletter et al. 2018). Similar to the case of Rbfox1, we observed that the highest percentage of muscle-expressed putative *mir-33* target genes are found in Mfuzz Clusters 35 and 36 (Supplemental Fig. S1C,D). To further explore the relationship between Rbfox1 and *mir-33*, we checked the overlap between the datasets of putative *mir-33* targets and Rbfox1 targets in each gene cluster (Supplemental Fig. S1E). Bolstering our claim, the maximum representation of common target genes, in terms of both absolute numbers and percentage of total, was found in Cluster 36, which contains *Rbfox1* (Supplemental Fig. S1E-H). Statistical analysis revealed a positive correlation between *mir-33* and Rbfox1 targets (Supplemental Fig. S1I,J). To conclude, given their antagonistic functions, our bioinformatic analysis suggests that Rbfox1 and *mir-33* may act at particular developmental time points to temper or fine-tune the expression of their common target genes during for muscle development.

Given the positive correlation between their respective target gene sets, we hypothesised that the expression profiles of Rbfox1 and *mir-33* may themselves be linked. Scouring their promoter regions, we found that the respective 1 kb-long promoter regions of Rbfox1 and *mir-33* share the binding sites of twenty-seven transcriptional regulators (Supplemental Fig. S2A,B). GO term analysis using AmiGO 2 (Carbon et al. 2009) revealed enrichment of the related biological processes of ‘BMP signaling pathway’, ‘activin receptor signaling pathway’, and ‘SMAD protein signal transduction’, which include the Dpp/BMP signalling components: MAD homology domain transcription factors, Mothers against dpp (Mad), Medea (Med), and Smad on X (Smox), and the C2H2 zinc finger transcription factor Schnurri (Shn) (Supplemental Fig. S2C). Similar analysis of the respective 5 kb-long promoter regions further yielded other components of the same signalling pathway, Daughters against dpp (Dad) and Sno oncogene (Snoo) (Supplemental Fig. S2D,E). Therefore, as per the bioinformatics data, Dpp/BMP signalling may lie upstream of both Rbfox1 and *mir-33.* However, further studies are needed to validate this finding.

### Core components of the JAK/STAT pathway are all involved in early adult myogenesis

Having established that *Stat92E* is critical for DLM template splitting, we embarked upon the systematic interrogation of the requirement of the other members of the JAK/STAT pathway in the same. Analysis of the transcriptomics resource of developing flight muscles, (Spletter et al. 2018), revealed that both *dome* and *hop* belong to Mfuzz Cluster 8, characterised by elevated expression from the myoblast stage until 30 h APF, followed by diminished expression until the mature adult muscle stage (Fig. 4A,B). Both *dome* and *hop* do not undergo alternative splicing, and did not appear in our bioinformatic survey of muscle genes with putative Rbfox1 binding motifs (Nikonova et al 2022).

**Figure 4.**
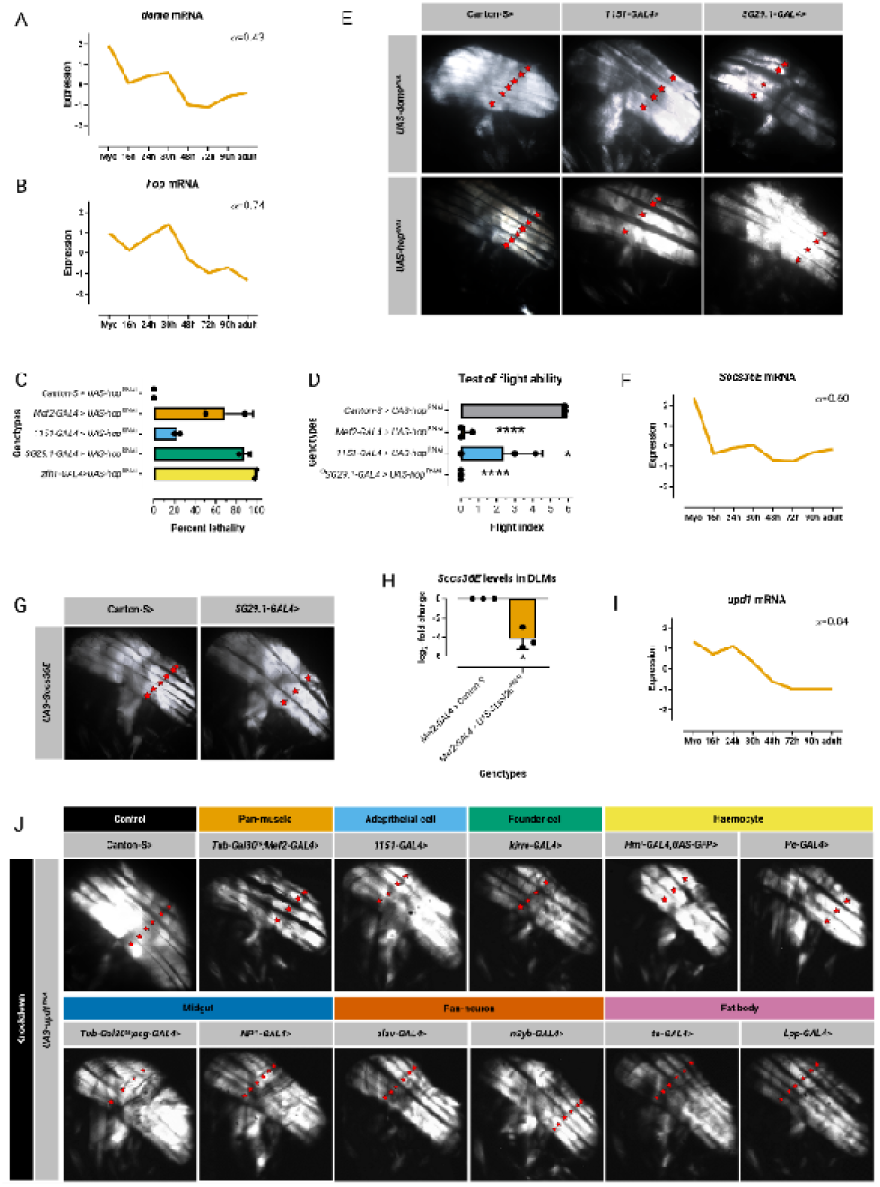
Core components of the JAK/STAT pathway are all involved in early adult myogenesis. Standard normal count values for *dome* **(A)**, *hop* **(B)**, *Socs36E* **(F)**, and *upd1* **(I)** from an mRNA-seq developmental time-course of wild-type IFMs (Spletter et al., 2018). **(C)** Quantification of pupal lethality from *hop* knockdown. Genotypes as noted. **(D)** Quantification of flight ability after *hop* knockdown. Genotypes as noted. Significance is from paired *t* test (\**P* < 0.05; \*\*\*\**P* < 0.0001). **(E, G, J)** Polarised microscopy images of hemithorax from flies. Genotypes as noted. Red stars indicate DLM fascicles. **(H)** Quantification of RT-qPCR data for *Socs36E* transcript levels in IFMs from *Stat92E* knockdown. Significance is from paired t test (\**P* < 0.05).

Because of their high expression at early myogenesis stage, we decided to study the effects of knockdown of *dome* and of *hop.* We observed that knockdown of *hop* causes pupal lethality (Fig. 4C), reminiscent of that of *Rbfox1* or *Stat92E* (Nikonova et al. 2022; Schnorrer et al. 2010). Moreover, knockdown of *hop* also significantly compromises flight ability (Fig. 4D). Muscle birefringence revealed that hemithoraces of flies with ‘early’ knockdown of either *dome* or *hop* had an abnormal number of four DLM fascicles (Fig. 4E). Together, we infer that the receptor Dome and the JAK Hop, core components of the JAK/STAT signalling pathway upstream to Stat92E, are also required for template splitting.

Next, we asked whether the cytokine ligands of the JAK/STAT signalling pathway, the three Upd glycoproteins, are required for DLM patterning too. Analysis of the transcriptomics resource of developing *Drosophila* flight muscles (Spletter et al. 2018), revealed that both *upd2* and *upd3* belong to Mfuzz Cluster 20, characterised by elevated expression at the myoblast stage, followed by sharp decline by 16 h APF, remaining low in expression until the adult muscle stage (Supplemental Fig. S5B,C). On the other hand, *upd1* belongs to Cluster 33, characterised by elevated expression from the myoblast stage until 24 h APF, followed by a gradual decline in expression until the adult muscle stage (Fig. 4I).

Because the Upd proteins are secreted glycoproteins that can act at a distance as the primary ligands for the JAK/STAT signalling pathway, to identify their source during early myogenesis, we used a battery of drivers, specific to, besides the muscle, the tissues adjacent to it. Muscle birefringence revealed that hemithoraces of flies with double knockdown of *upd2* and *upd3* had normal number of six DLM fascicles (Supplemental Fig. S5F), suggesting that, consistent with their low expression during early myogenesis stage, the ligands are dispensable for template splitting. On the other hand, muscle birefringence revealed that hemithoraces of flies with knockdown of *upd1* in muscle, haemocytes, and adult midgut precursors had an abnormal number of four DLM fascicles (Fig. 4J), suggesting that Upd1 is required both cell-autononously and non-cell-autononously for template splitting.

### Rbfox1 putatively destabilises the *Socs36E* transcript, which encodes the JAK/STAT signalling pathway inhibitor

The *Suppressor of cytokine signaling at 36E* (*Socs36E*) gene, which is a target of the JAK/STAT pathway, encodes inhibitory proteins that promote degradation of Dome and Hop, thereby providing a negative-feedback loop (Stec et al. 2013). As per the transcriptomics resource of developing flight muscles (Spletter et al. 2018), *Socs36E*, like *upd2* and *upd3*, belongs to Mfuzz Cluster 20 (Fig. 4F). Therefore, we hypothesised whether its low expression by 16 h APF is required for Stat92E activity to peak at this time point. To determine whether *Socs36E* is a direct negative regulator of JAK/STAT signalling during early myogenesis, we overexpressed *Socs36E* driven by *SG29.1-GAL4*, and muscle birefringence revealed that the hemithoraces of progeny flies had an abnormal number of three DLM fascicles (Fig. 4G).

Finally, we asked, given that *Socs36E* is a Stat92E target gene in other tissues, why its mRNA expression remain low when Stat92E is active. We tested two possibilities. First, at the transcriptional level, perhaps Stat92E does not regulate *Socs36E* expression in the developing flight muscles. Contrary to this hypothesis, we found that *Socs36E* transcripts are significantly downregulated in the genetic background of *Stat92E* knockdown (Fig. 4H), confirming that Stat92E-mediates transcription of *Socs36E* during myogenesis. Second, maybe *Socs36E* transcripts are negatively regulated at the post-transcriptional level.

Interestingly, *Socs36E* mRNA bears a *GCAUG* motif within its short, 858 bases-long 3’-UTR (Supplemental Fig. S3G). It has been shown that, in the *Drosophila* ovary, the presence of *GCAUG* elements within the 3’-UTR of the *pumilio* mRNA results in decreased stability (Carreira-Rosario et al. 2016). Therefore, we postulate that, despite high Stat92E expression during early adult myogenesis stage, the coincidentally high levels of Rbfox1 help destabilise the *Socs36E* mRNA, repressing their translation, and preventing Socs36E from interfering with the activity of the JAK/STAT pathway. On the other hand, declining Rbfox1 expression, during mid adult myogenesis stage, frees the lingering *Socs36E* mRNA from repression, allowing the protein product to inhibit the JAK/STAT signalling pathway in the developing DLMs at that time point.

### Stat92E and Rbfox1 together activate the stemness regulator Zfh1 during myogenesis

Although JAK/STAT signalling also controls homoeostatic functions such as proliferation and survival, the direct transcriptional STAT targets that regulate these processes are largely unknown. Through its role in certain *Drosophila* tissues, some direct targets of JAK/STAT signalling involved in various axes of homoeostasis and regeneration have been identified: Stat92E influences F-actin dynamics in germline stem cells through the *Drosophila* profilin orthologue Chickadee (Chic), and facilitates GSC-niche attachment (Shields et al. 2014); the anti-apoptotic factor Death-associated inhibitor of apoptosis 1 (Diap1) acts downstream of Stat92E in maintaining the viability of posterior wing cells during development (Recasens-Alvarez et al. 2017); and Zfh1 promotes CySC self-renewal (Leatherman and DiNardo 2008).

Zfh1 encodes a transcriptional repressor required for numerous mesodermal cell fate decisions during embryogenesis (Su et al. 1999). To explore its role during early myogenesis, we undertook knock-down experiments of *zfh1*. Flies with ‘early’ knock-down of *zfh1* exhibited impaired flight ability (Fig. 5B), and had an abnormal number of four DLM fascicles per hemithorax (Fig. 5D). Previous studies have established that Zfh1 plays a central role in maintaining the FCMs in the undifferentiated state until fusion by inhibiting the myogenic signal provided by Mef2 function (Boukhatmi and Bray 2018). Consequently, Zfh1-positive cells are the actively dividing myoblasts that swarm the persistent larval muscles. We suspect that knock-down of *zfh1* during the early adult myogenesis stage is likely to precociously imbue the myoblasts with myogenic potential, and hence can result in fewer cells in the FCM pool. Our lab has previously shown that an optimum number of myoblasts is required for achieving the normal IFM size and pattern (Rai and Nongthomba 2013). Hence, a decreased number of actively dividing myoblasts may be the reason behind the splitting defects revealed by our work on knock-down of *zfh1*.

**Figure 5.**
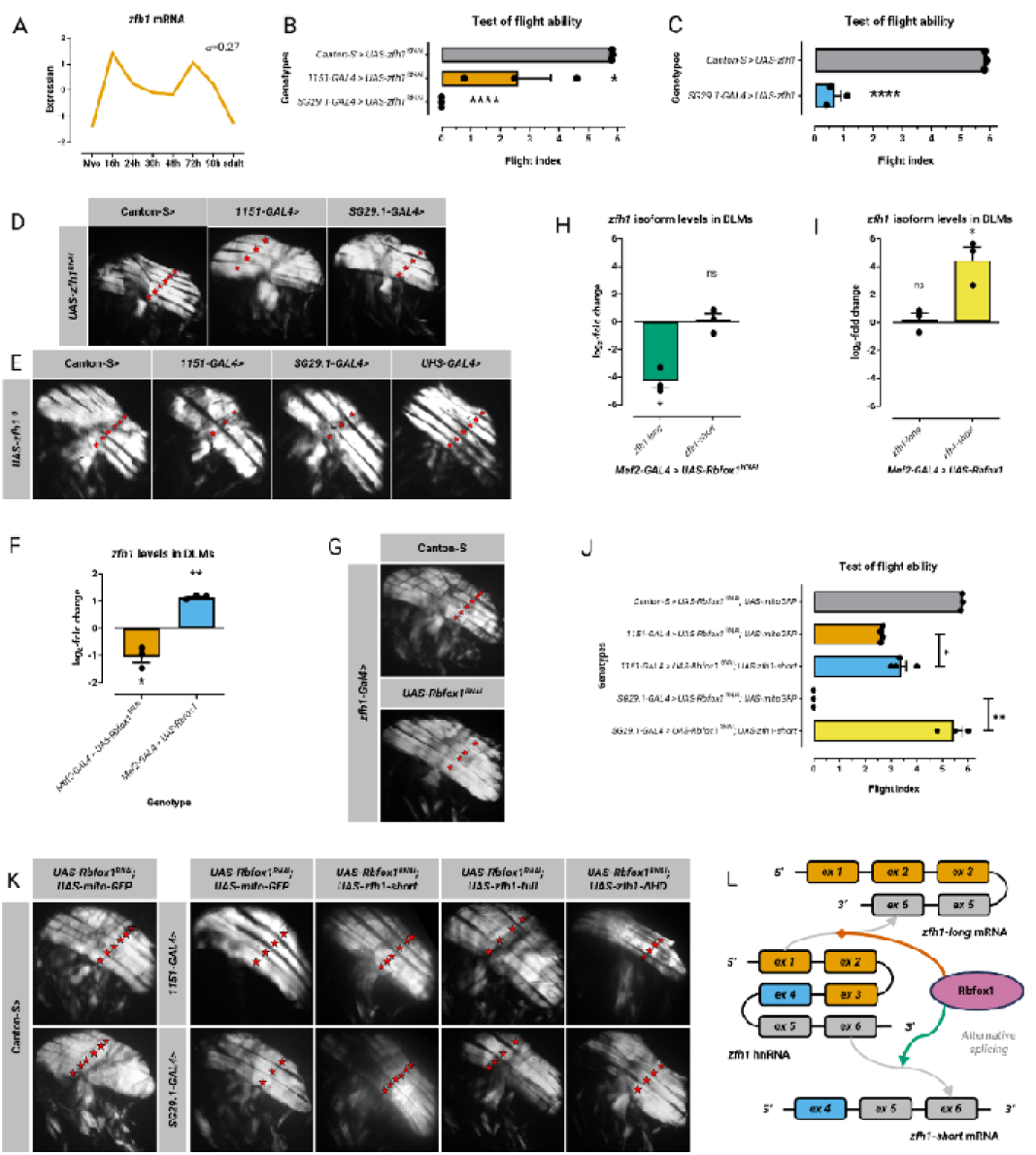
Rbfox1 regulates the alternative splicing of *zfh1* which is necessary for splitting of the DLM templates. **(A)** Standard normal count values for *zfh1* from an mRNA-seq developmental time-course of wild-type IFMs (Spletter et al., 2018). Quantification of flight ability after *zfh1* knockdown **(B)** and overexpression **(C)**. Genotypes as noted. Significance is from paired *t* test (\**P* < 0.05; \*\*\*\**P* < 0.0001). **(D, E, G, K)** Polarised microscopy images of hemithorax from flies. Genotypes as noted. Red stars indicate DLM fascicles. **(F)** Quantification of RT-qPCR data for *zfh1* transcript levels in IFMs from *Rbfox1* misexpression. Significance is from paired t test (\**P* < 0.05; \*\**P* < 0.01). Quantification of RT-qPCR data for *zfh1* splice variant levels in IFMs from *Rbfox1* knockdown **(H)** and overexpression **(I)**. Significance is from paired t test (\**P* < 0.05). **(J)** Quantification of flight ability after *zfh1* overexpression in *Rbfox1* knockdown Genotypes as noted. Significance is from paired *t* test (\**P* < 0.05; \*\*\*\**P* < 0.0001). **(L)**. Graphical summary.

*Rbfox1*, *Stat92E*, and *zfh1* share the same temporal expression profile (Fig. 5C). Indeed, *zfh1* mRNA bears multiple Rbfox1 binding sites (Supplemental Fig. S3H). To establish whether Rbfox1 regulates *zfh1*, we interrogated whether *Rbfox1* misexpression affects the transcript levels of *zfh1*. We found that that there is a direct correlation between *Rbfox1* expression and *zfh1* expression: knock-down of *Rbfox1* results in downregulation of *zfh1*, whereas overexpression of *Rbfox1* causes upregulation of *zfh1* (Fig. 5F). Indeed, we found that even *zfh1-GAL4-*driven knock-down of *Rbfox1* perturbs splitting of the DLM templates, suggesting the cell-autonomous requirement of Rbfox1 for *zfh1* expression (Fig. 5G).

Development requires optimum expression levels, and we have reported how overexpression of *Rbfox1*, like its knockdown, is catastrophic for myogenesis (Nikonova et al. 2022). Similar to that observation, we found that overexpression of *zfh1* is pupal lethal. Driving the overexpression by *SG29.1-GAL4*, at milder 25°C, produced offspring with impaired flight ability (Fig. 5C). Moreover, muscle birefringence revealed that overexpression of *zfh1* completely blocks template splitting, and, so, hemithoraces of flies had only three DLM fascicle (Fig. 5E). Thus, we infer that the expression of *Rbfox1* and its targets must be tightly regulated at different developmental time points for myogenesis to proceed properly, and we have indeed discovered such a mechanism (A Mukherjee and U Nongthomba, under revision).

### *Misregulation of* Rbfox1 *has isoform-specific effects on* zfh1 *expression*

*zfh1* exists in two isoforms, a long (*zfh1-long*) and a short one (*zfh1-short*) (Boukhatmi and Bray 2018). We found multiple intronic (*U)GCAUG* sites directly upstream of the alternative exon, which is retained in *zfh1-short* but is excluded from *zfh1-long* (Fig. 5L). We hypothesised that Rbfox1 regulates the alternative splicing of *zfh1*, and to verify this, we decided to interrogate the expression levels of the *zfh1* isoforms in the genetic backgrounds of *Rbfox1* misexpression. We found that in case of *Rbfox1* knock-down the expression of the *zfh1-long* transcript is downregulated, whereas there is no significant change in the expression of the *zfh1-short* transcript (Fig. 5H). This suggests that the downregulation of total *zfh1* expression observed in case of *Rbfox1* knock-down is contributed to by the repression of the *zfh1-long* isoform alone. On the other hand, *Rbfox1* overexpression caused an upregulation in the expression of only the *zfh1-short* transcript, but did not affect the expression of the *zfh1-long* counterpart (Fig. 5I). This, in turn, indicates that the total *zfh1* upregulation observed in case of *Rbfox1* overexpression reflected the upregulation of the *zfh1-short* transcript alone. Collectively, these results demonstrate that Rbfox1 is involved in the alternative splicing of the *zfh1* transcript: the advent of *Rbfox1* expression during early adult myogenesis perhaps marks a shift in the dynamics of *zfh1* isoforms, with a switch from the default *zfh1-long*, expressed in all muscle progenitors at larval stage, to the *zfh1-short*, which is maintained in pupal muscle progenitors (Fig. 5L). Thus, this finding adds a new layer of complexity to the preferential regulation of a specific splice variant of *zfh1*.

To test this genetic interaction, we directed the ‘early’ overexpression of *UAS-zfh1*, *UAS-zfh1-short*, and *UAS-zfh1-*Δ*HD* in the genetic background *Rbfox1* knock-down. Polarised light microscopy showed that concomitant overexpression of *UAS-zfh1-short* is able to suppress the *Rbfox1* knock-down phenotype: six DLM fascicles were observed per hemithorax (Fig. 5K). On the other hand, concomitant overexpression of *UAS-zfh1-*Δ*HD* fails to suppress the *Rbfox1* knock-down phenotype (Fig. 5K), suggesting that its homoeodomain is essential for Zfh1 function during myogenesis. Even the full-length *zfh1* cDNA is insufficient (Fig. 5K) because, due to the dearth of Rbfox1, it cannot be spliced to generate the functional *zfh1-short* isoform. Interestingly, reflecting the rescue of the DLM splitting defect, *UAS-zfh1-short* also significantly improves the flight ability of *Rbfox1^RNAi^* flies (Fig. 5J). Together, we infer that Rbfox1 is required to catalyse the formation of Zfh1-short, and its DNA-binding activity is critical for DLM biogenesis.

Similar to our analysis of the Stat92E isoforms, we predicted the structural differences between the Zfh1 isoforms encoded by the splice variants under native conditions. We found that the long (Zfh1-PB) isoform harbours an additional C2H2-type zinc finger domain (Pfam entry: PF13912) at its N-terminus, which is absent in the short isoform (Zfh1-PA) (Supplemental Fig. S6A,B). Although C2H2 zinc finger motifs are classically known to recognise DNA sequences, they can also bind to RNA and protein targets (Wolfe et al. 2000). Furthermore, the long isoform has a higher score LLR, suggesting that it is more disordered than the short isoform. Unique to itself, the long isoform bears two PrLDs between residues 100-300 (Supplemental Fig. S6C,D). PrLDs promote multivalent and transient interactions involving RNA, structured domains, and PrLDs of the same and/or other proteins (Malinovska et al. 2013) Based on these data, we propose that the Zfh1 isoforms have overlapping but also distinct functions: the long isoform, aided by its additional C2H2-type zinc finger domain, may engage in a unique repertoire of promiscuous interactions, compared to the short isoform.

### *The Stat92E downstream effector gene* Diap1 *prevents apoptosis of migrating myoblasts*

Studies in imaginal discs have shown that activated Stat92E can directly increase Diap1, an E3 ubiquitin ligase with a caspase inhibitor activity, which prevents apoptosis after a variety of stresses (Betz et al. 2008). *Mef2-GAL4*-driven RNAi of *Diap1* resulted in late larval lethality (Schnorrer et al. 2010). As per the transcriptomics resource of developing flight muscles (Spletter et al. 2018), *Diap1* belongs to Cluster 20, characterised by its highest expression at the myoblast stage, and gradual decline by 16 h APF (Fig. 6A). Interestingly, there are multiple intronic Rbfox binding sites flanking the five alternative exons of *Diap1* (Supplemental Fig. S3I), suggesting that it is both directly and indirectly regulated by Rbfox1, similar to some other Stat92E targets, *Socs36E*, *zfh1*, and *chic*.

**Figure 6.**
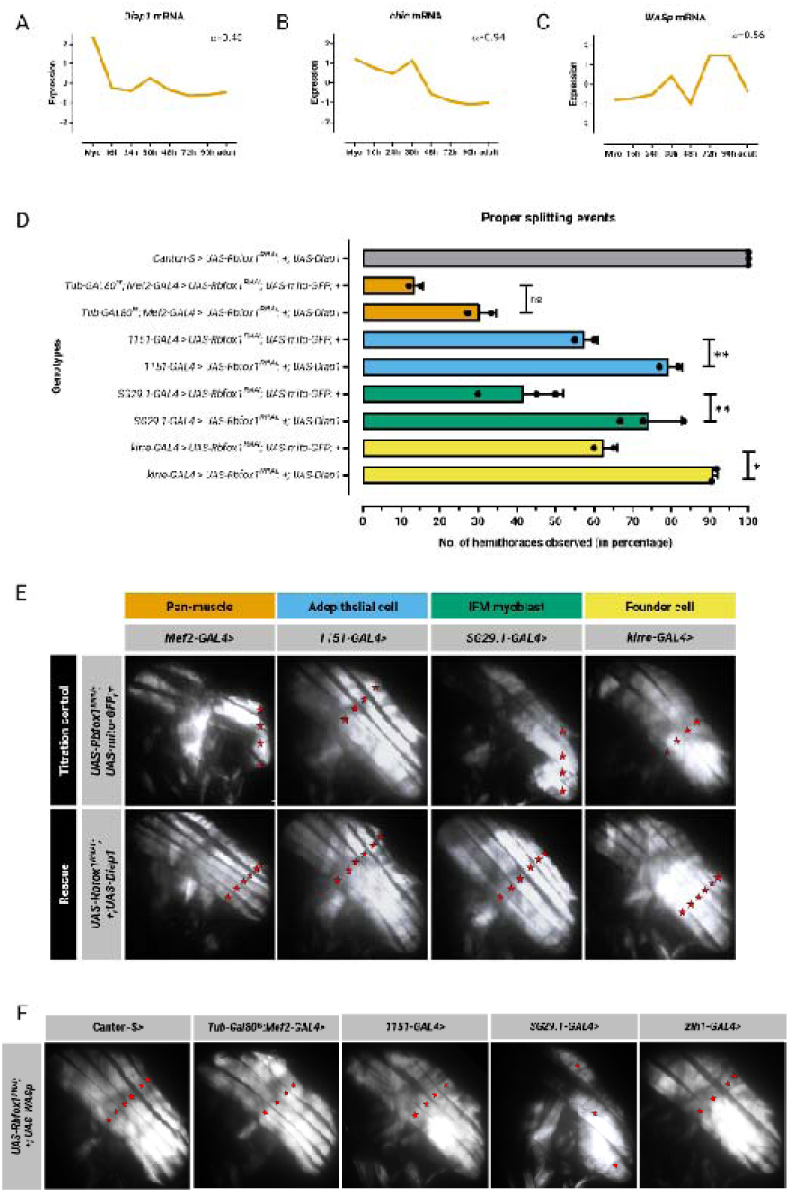
Stat92E targets Chic and Diap1 are necessary for splitting of the DLM templates. Standard normal count values for *Diap1* **(A)**, *chic* **(B)**, and *WASp* **(C)** from an mRNA-seq developmental time-course of wild-type IFMs (Spletter et al., 2018). **(D)** Quantification of flight ability after expression of *Diap1* in the genetic background of *Rbfox1* knockdown. Genotypes as noted. Significance is from paired *t* test (\**P* < 0.05; \*\**P* < 0.01). **(E, F)** Polarised microscopy images of hemithorax from flies. Genotypes as noted. Red stars indicate DLM fascicles.

To test our hypothesis, we directed the overexpression of *Diap1*, in the genetic background of *Rbfox1* knock-down. Expression of *UAS-Diap1* significantly improves the flight ability of *Rbfox1^RNAi^* flies (Fig. 6D). Polarised light microscopy showed that concomitant overexpression of *Diap1* can suppress the *Rbfox1* knock-down phenotype: six DLM fascicles were observed per hemithorax (Fig. 6E). Based on these results, we infer that *Diap1* lies downstream of Rbfox1 during DLM biogenesis, and allows migrating myoblasts to survive.

### *Rbfox1 putatively regulates* ena *and the Stat92E target gene* chic *to facilitates myoblast fusion to the larval muscle templates*

Compared to the four Profilin genes identified in mammals, *Drosophila* has a single Profilin orthologue, encoded by *chic* (Cooley et al. 1992). Profilins promote actin polymerisation by catalysing ADP to ATP exchange on globular actin, and through transient interactions of the profilin–ATP–actin complex with the fast-growing ‘barbed’ end of F-actin (Kooij et al. 2016).

Moreover, Profilin contains a phosphatidylinositol 4,5-bisphosphate binding motif, and interacts with phosphatidylinositol lipids and transcription factors, highlighting its roles in signalling and gene activity (Kooij et al. 2016; Staudt et al. 2005). During the fusion of FCMs with the larval muscle templates, multiple long actin-based protrusions, called filopodia, emanate from the template surface, and are necessary for template-myoblast adhesion (Segal et al. 2016). Chic influences filopodia-forming processes at different steps, and stimulates the anti-capping activities of Enabled (Ena) (Gonçalves-Pimentel et al. 2011). The knockdown of *ena* under control of the myotube-specific driver *kirre^rP298^-GAL4* (Menon and Chia 2001) gives rise to defects in template splitting (Segal et al. 2016). Interestingly, *ena* too undergoes alternative splitting, and bears multiple Rbfox1 binding sites (Supplemental Fig. S3K).

Analysis of the transcriptomics resource of developing flight muscles (Spletter et al. 2018) revealed that *chic*, similar to *dome* and *hop*, belongs to Cluster 8 (Fig. 6B). A few systematic screens have identified the effect of *chic* on muscle morphogenesis and function in *Drosophila*: a gain-of-function screen for genes that affect embryonic muscle pattern formation showed that *sr-GAL4*-driven overexpression of *chic* caused variable weak to extreme phenotypes (Staudt et al. 2005), whereas *Mef2-GAL4*-driven RNAi of *chic* resulted in embryonic lethality (Schnorrer et al. 2010). *chic* bears, at its 5’ end, an Rbfox1 binding site between its alternative exons and its constitutive exon (Supplemental Fig. S3J).

Thus, given that fusion is blocked when formation of these filopodia is compromised, we hypothesise that the downregulation of chic, in the templates, in the genetic background of *Stat92E* knockdown, affects filopodia formation, and, consequently, myoblast fusion, resulting in the patterning phenotype. To validate whether it is indeed the deficiency of *chic* in the templates which is responsible for the splitting defect, we decided to adopt the corollary approach by overexpressing a factor promoting actin-polymerisation in the apposing FCM. We chose WASp (Fig. 6C), the *Drosophila* orthologue of the conserved Wiskott-Aldrich Syndrome proteins, which is required, specifically in the FCMs, for the formation of the F-actin foci, and promoting efficient foci invasion into the apposing FC with multiple finger-like protrusions, called podosomes, which evolve into single-channel fusion pores between the two muscle cells (Sens et al. 2010). Overexpression of *WASp*, in the adepithelial cells using *1151-GAL4*, and in the IFM myoblasts using *SG29.1-GAL4*, in the genetic background of *Rbfox1* knockdown, revealed that hemithoraces of the progeny had an abnormal number of four or three DLM fascicles, respectively (Fig. 6F). This suggests that increasing the expression level of WASp is not sufficient rectify the *Rbfox1* knockdown phenotype because the fusion defect does not lie at FCM side, but is instead particular to the templates. Thus, knockdown of *Rbfox1* and of *Stat92E*, via the downregulation of *chic* in the templates, may result in filopodia formation defects, which compromise myoblast fusion.

## Discussion

### Pleiotropic roles of Rbfox1 during myogenesis

In this study, we have identified multiple roles of Rbfox1 during DLM development, ranging from maintaining stemness and survival of myoblasts, to promoting their eventual fusion with the templates (Graphical summary). To uncover other putative direct targets of Rbfox1 during flight muscle development, we superimposed the genes with Rbfox1 motif instances (Nikonova et al. 2022), identified using oRNAment (Bouvrette et al. 2020), against genes representing GO terms for various biological processes. Interestingly, we found *svp* and *Stat92E* in the overlap with the GO biological process terms ‘cell population proliferation’ and ‘regulation of cell population proliferation’, respectively (Supplemental Fig. S7A,B). The overlap with the GO biological process term ‘regulation of programmed cell death’ revealed *Deubiquitinating apoptotic inhibitor* (*DUBAI*), which encodes a cysteine-type deubiquitinase involved in negative regulation of apoptosis (Supplemental Fig. S7C,D). DUBAI physically interacts with Diap1, stabilising it, despite presence of death-inducing stimuli that would induce Diap1 degradation (Yang et al. 2014). We hypothesise that Rbfox1-targetted activation of DUBAI maintains the abundance of Diap1, even though its mRNA expression declines by 16 h APF (Spletter et al. 2018; Fig. 6C). Finally, the overlap with the GO biological process term ‘actin filament organization’ revealed *Gelsolin* (*Gel*) and *CG43901* (Supplemental Fig. S7E,F). There is scant information available on the product of *CG43901*: while it lacks conserved domains, it features disordered regions, and is predicted to be cytoplasmic. On the other hand, Gel is a well-characterised actin interactor belonging to the conserved Gelsolin/Villin family, and has been shown, by limiting myoblast fusion, to regulate the size and splitting of a subset of larval body wall muscles, the lateral transverse muscles (Bertin et al. 2021). Future studies may further investigate the role of these and other Rbfox1 targets in muscle development.

### *Rbfox1 and* mir-33 *function appears to be coupled across evolution*

Our study has implications for areas as distant as evolution and pathology. We have found that antagonism between Rbfox1 and *mir-33* tempers gene expression of their common target mRNA, *Stat92E.* Moreover, our bioinformatic analysis hints that the overlap between their targets may be more extensive. We have found instances of their involvement in common biological processes in the literature. Both RBFOX1 activity, and *miR-33* expression are reduced in skeletal muscle biopsies from myotonic dystrophy type-1 patients (Klinck et al. 2014; Perbellini et al. 2011; reviewed in Mukherjee and Nongthomba 2023). Data analysis of a genome-wide association study has implicated variations within the *RBFOX1* locus with influencing obesity (Ma et al. 2010; reviewed in Mukherjee and Nongthomba 2023). Similarly, *MIR33A*, located within intron 16 of the gene encoding Sterol regulatory element binding transcription factor 2 (SREBF2), a key transcriptional regulator of cholesterol synthesis, is causal to both Abdominal obesity-metabolic syndrome 1 and Non-alcoholic fatty liver disease (Marquart et al., 2010; Rayner et al. 2010). *mir-33* has an analogous role in *Drosophila*, ensuring balanced lipid metabolism by targeting genes involved in both breakdown and formation of fatty acids and triacylglycerol (Clerbaux et al. 2021). It has been shown that the skeletal muscle communicates with other organs to prevent excess fat storage, and thereby protects against obesity (Eckel 2019; Ghosh et al. 2020), and that *mir-33* serves to prevent extreme fluctuations in metabolically sensitive tissues (Clerbaux et al. 2021). Therefore, we propose that metabolism may play a role in the development of striated muscles, with Rbfox1 and *mir-33* being key players in that conserved crosstalk. It may be interesting to study how this interaction affects the allocation of energy stores to adult myogenesis considering that, since pupae do not feed, metamorphosis is primarily fuelled by lipid catabolism (Merkey et al. 2011).

### A novel function of the JAK/STAT pathway during myogenesis

Grasping how cells communicate during development not only provide insights into mechanisms of cell communication in organ formation, but also a clearer picture of why congenital defects occur. Hints into the role of the JAK/STAT signalling pathway in muscle development may be gleaned indirectly. Mice deficient in the *suppressor of cytokine signalling 2* (*SOCS2*), a human orthologue of *Socs36E*, have increased muscle cell numbers (Metcalf et al. 2000). Failure of destabilisation and ubiquitination of Protein inhibitor of activated STAT 4 (PIAS4) has been associated with Limb-girdle muscular dystrophy type 2H (Albor et al. 2006). Moreover, interestingly, JAKs are not the only upstream kinases of STATs: activated MET proto-oncogene, receptor tyrosine kinase (MET) is able to recruit STAT3 among several other signalling effectors, inducing its phosphorylation, triggering its dimerisation and nuclear translocation (Boccaccio et al. 1998). Remarkably, mutation in the *MET* gene has been implicated in skeletal muscle dysplasia, and demonstrated to be crucial for the migration of muscle progenitor cells, and the proliferation of secondary myoblasts in mice (Zhou et al. 2019). Interestingly, when we interrogated the time point-specific transcriptomes of developing Drosophila flight muscles (Spletter et al. 2018), we found that the *Drosophila* orthologue of *MET*, *Ror* mRNA has elevated expression from the myoblast stage until 24 h APF, with a peak in mRNA expression at 16 h APF (Supplemental Fig. S3B), which suggests that Ror may have a role in early adult myogenesis. Thus, our study provides a suitable entry point to explore the question of JAK/STAT signalling pathway in myogenesis, having provided fundamental insights into how knockdown of *Stat92E*, and the overexpression of *Socs36E* prevent DLM template splitting, which is known to be affected by myoblast numbers (Rai and Nongthomba 2013). Future studies may explore Ror is required for the activation of Stat92E, perhaps redundantly with the *Drosophila* JAK, Hop.

*JAK* mutations and genetic alterations in downstream components of the JAK/STAT pathway, including mutations in *STAT* genes, are very frequent in haematological malignancies, disorders that arise from the haematopoietic stem cells (Vainchenker and Constantinescu 2013). Interestingly, the binding of *Mir33* to two conserved motifs in the 3’-UTR of *transformation related protein 53*, which encodes tumour protein p53, controls haematopoietic stem cell self-renewal in mouse (Herrera-Merchan et al. 2010). This suggests that the *mir-33-*mediated regulation of *Stat92E*, which we have discovered in this study, may be universal, prevailing in contexts outside of myogenesis too.

## Materials and methods

### Fly stocks

Canton-S was used as the wild-type strain. The transgenic fly lines used were procured from Bloomington Drosophila Stock Center, Indiana University, while the following fly stocks were kind gifts: *UAS-mir-33*, *UAS-mir-33.sponge*, and *UAS-scramble.sponge* from Dr Jennifer Kennell (Vassar College, Poughkeepsie, United States of America); *UAS-zfh1*, *UAS-zfh1-short*, and *UAS-zfh1-*Δ*HD* from Dr Hadi Boukhatmi (IGDR, Rennes, France); *UAS-zfh1^RNAi^* and *zfh1-GAL4* from Prof K. VijayRaghavan (NCBS, Bangalore, India); *UAS-Rbfox1* and *UAS-Rbfox1^RNAi^* from Prof L. S. Shashidhara (IISER, Pune, India); *UAS-Diap1* from Prof Girish Ratnaparkhi (IISER, Pune, India); *UAS-upd1^RNAi^* and *UAS-Socs36E* from Dr Tina Mukherjee (InStem, Bangalore, India); and *He-GAL4*, *Hml-GAL4,UAS-GFP*, *Tub-GAL80^ts^,esg-GAL4*, *NP1-GAL4*, *UAS-upd2^RNAi^;UAS-upd3^RNAi^*, and *UAS-Stat92E^RNAi^* from Dr Sveta Chakrabarti (IISc, Bangalore, India).

### Behavioural assays

Flies were aged for 2-3 days before testing for flight ability, and the assay was performed as described by Drummond et al., (1991). The flight ability data were then transformed into a flight index for each genotype, determined by using, with slight modifications, the formula described by Tohtong et al. (1995): 6 U/T + 4 H/T + 2 D/T + 0 F/T, where U (up), H (horizontal), D (down), and F (flightless) are the number of flies in each category of flight ability, and T is the total number of flies tested for that genotype.

### Polarised microscopy

Sample preparation for imaging the IFMs in the adults was done as described by Nongthomba & Ramachandra (1999). The hemithorax mounts were prepared using DPX mountant (SDFCL), and observed under an Olympus SZX12 microscope, and photographed using an Olympus C-5060 camera under polarised light optics.

### Real-time quantitative reverse transcription PCR (RT-qPCR)

Quantitative PCR was carried out using HOT FIREPol EvaGreen qPCR Supermix (5X, Solis BioDyne). *Ribosomal protein L32* (*RpL32*, previously known as *rp49*) served as an internal control for normalisation in all reactions. The 2^-ΔΔC^_T_ method was used as the relative quantification strategy the qPCR data analysis (Livak & Schmittgen, 2001). All primers are listed in Supplemental Table S1.

### Bioinformatic analysis

FlyBase (release FB2023_06) was used to find information on phenotypes, function, stocks, gene expression, etc. Functional classification of gene lists, and expression data files were performed using the PANTHER Gene List Analysis tool (Thomas et al., 2022). Gene Ontology term enrichment for biological process was performed using the AmiGO 2 Term Enrichment Service (Carbon et al., 2009). Rbfox1 motif and *mir-33* target site instances at regions of interest within the *D. melanogaster* (Aug 2014 BDGP Rel6 + ISO1 MT/dm6) genome assembly were bioinformatically identified using the PWMScan tool (Ambrosini et al., 2018). MicroRNA matches to *Rbfox1* and *Stat92E* 3’-UTRs, and predicted fly microRNA targets of the conserved microRNA family *mir-33-5p* were retrieved using TargetScanFly (Agarwal et al., 2018). The UCSC Table Browser tool (Karolchik et al., 2004) was used to retrieve and export ReMap 2022 (Hammal et al., 2022) track data for regions of interest. Protein disorder was predicted using the IUPred2A web interface (Erdős & Dosztányi, 2020), and the prediction algorithm PLAAC (Lancaster et al., 2014).

## Acknowledgments

We thank Vishakha Nesari, Upasana Gupta, and Sautan Show for helpful discussions. We acknowledge the researchers for generously sharing and the Bloomington Stock Center for reliable provision of the transgenic fly stocks used in this study. The work and A.M. were supported by the Indian Institute of Science, Bangalore [IE/REDA-23-1788-18], and the Council of Scientific and Industrial Research, New Delhi [09/079(2787)/2018-EMR-I], respectively.

