## Supplemental Figures for "Rbfox1 and *mir-33* regulate pleiotropic roles of JAK/STAT signalling during adult myogenesis in *Drosophila melanogaster*"

**Mukherjee_Graphical_Summary**

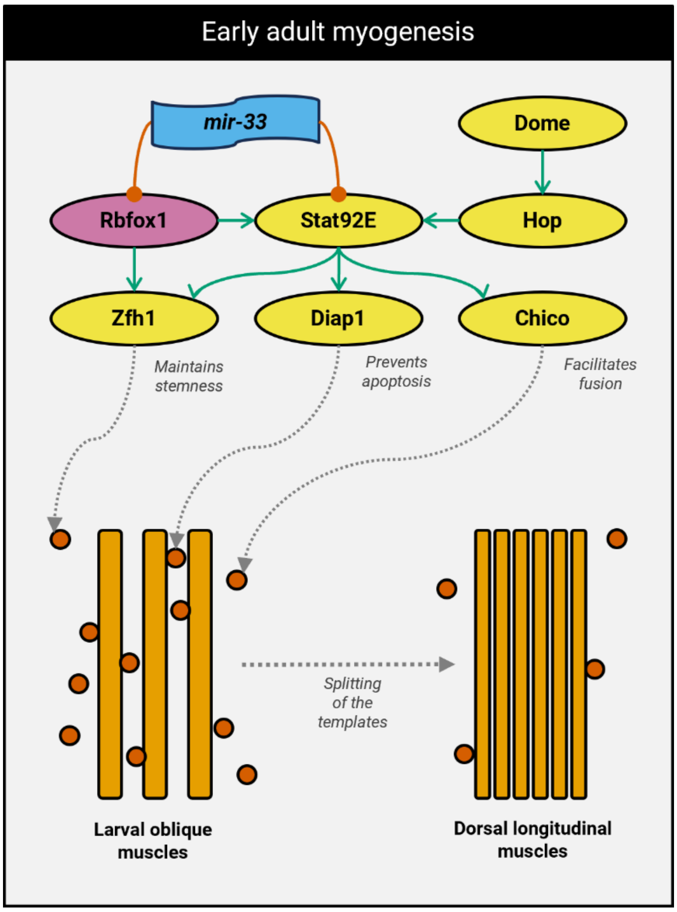


**Supplemental Figure. (Graphical Summary)** A model of the genetic interactions during early adult myogenesis in *Drosophila*.

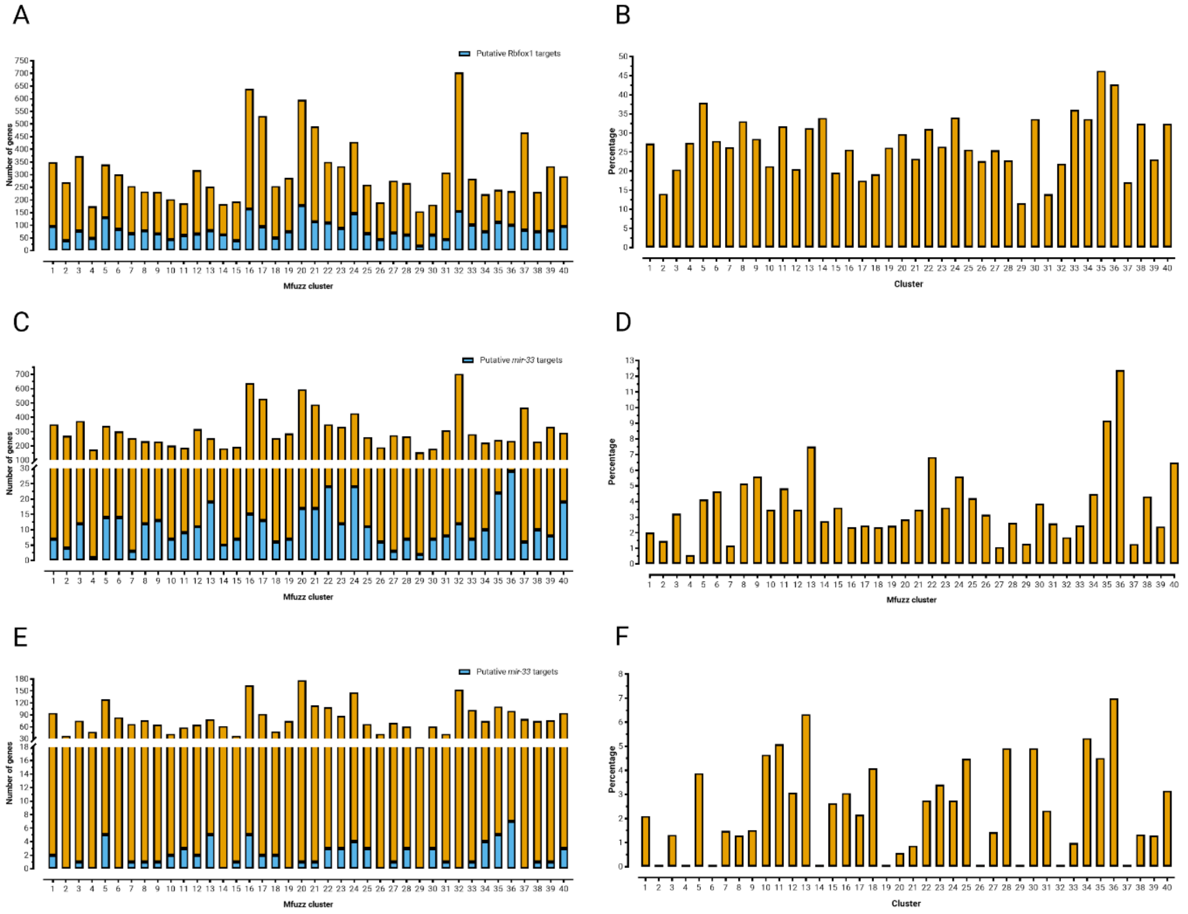


G

| FlyBase ID | Annotation Symbol | Symbol | Name | Gene Group (FlyBase) | Protein Signatures (InterPro) |
|---|---|---|---|---|---|
| FBgn0261871 | CG33507 | dpr2 | defective proboscis extension response 2 | DEFECTIVE PROBOSCIS EXTENSION RESPONSE | Immunoglobulin-like domain superfamily |
| FBgn0053516 | CG33516 | dpr3 | defective proboscis extension response 3 | DEFECTIVE PROBOSCIS EXTENSION RESPONSE | Immunoglobulin-like domain superfamily |
| FBgn0039862 | CG1804 | kek6 | kekkon 6 | KEKKONS | Immunoglobulin-like domain superfamily |
| FBgn0026206 | CG12218 | mei-P26 | meiotic P26 | TRIM UBIQUITIN LIGASES | Zinc finger, RING-type |
| FBgn0260660 | CG42543 | Mp | Multiplexin | COLLAGENS | Collagen type XV/XVIII, trimerization domain |
| FBgn0000036 | CG5610 | nAChRα1 | nicotinic Acetylcholine Receptor α1 | NICOTINIC ACETYLCHOLINE RECEPTOR SUBUNITS | Neurotransmitter-gated ion-channel |
| FBgn0016917 | CG4257 | Stat92E | Signal-transducer and activator of transcription protein at 92E | UNCLASSIFIED DNA BINDING DOMAIN TRANSCRIPTION FACTORS | p53-like transcription factor, DNA-binding domain superfamily |

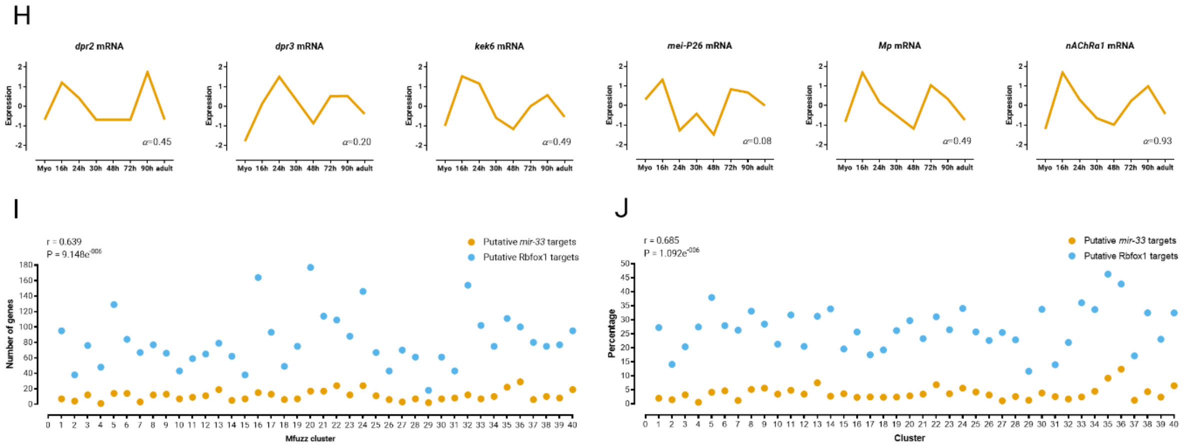


**Supplemental Figure 1.** Frequency distribution of number of putative Rbfox1 **(A)** and *mir-33* **(C)** targets (sky blue) out of total number of genes (orange) per Mfuzz Cluster. Frequency distribution of percentage of total number of genes that are putative Rbfox1 **(B)** and *mir-33* **(D)**

targets per Mfuzz Cluster. **(E)** Frequency distribution of number of putative *mir-33* targets (sky blue) out of total number of putative Rbfox1 target genes (orange) per Mfuzz Cluster. **(F)** Frequency distribution of percentage of total number of putative Rbfox1 target genes that are putative *mir-33* targets per Mfuzz Cluster. **(G)** List of putative shared targets of Rbfox1 and *mir-33* in Cluster 36, and **(H)** their standard normal count values from an mRNA-seq developmental time-course of wild-type IFMs (Spletter et al., 2018). Scatter plots of number **(I)** and percentage **(J)** of putative Rbfox1 (sky blue) and *mir-33* targets (orange) per Mfuzz Cluster to measure the correlation between them.

**Mukherjee_Supplemental_Fig_S2**

A D

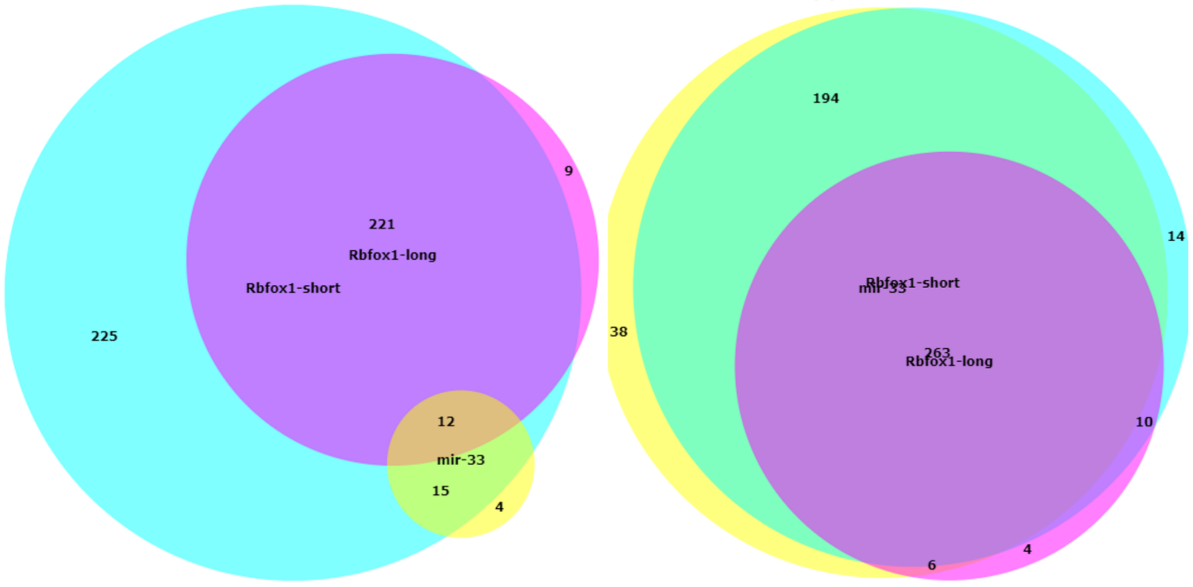


B

| FlyBase ID | Symbol |
|---|---|
| FBgn0087035 | AGO2 |
| FBgn0015602 | BEAF-32 |
| FBgn0262975 | cnc |
| FBgn0000283 | Cp190 |
| FBgn0000439 | Dfd |
| FBgn0000606 | eve |
| FBgn0004656 | fs(1)h |
| FBgn0014179 | gcm |
| FBgn0001168 | h |
| FBgn0041210 | HDAC4 |
| FBgn0039743 | Jasper |
| FBgn0002441 | l(3)mbt |
| FBgn0011648 | Mad |
| FBgn0011655 | Med |
| FBgn0014340 | mof |
| FBgn0005616 | msl-2 |
| FBgn0004861 | ph-d |
| FBgn0004860 | ph-p |
| FBgn0005624 | Psc |
| FBgn0003330 | Sce |
| FBgn0003396 | shn |
| FBgn0025800 | Smox |
| FBgn0010278 | Ssrp |
| FBgn0003567 | su(Hw) |
| FBgn0003607 | Su(var)205 |
| FBgn0036398 | upSET |
| FBgn0259789 | zld |

C E

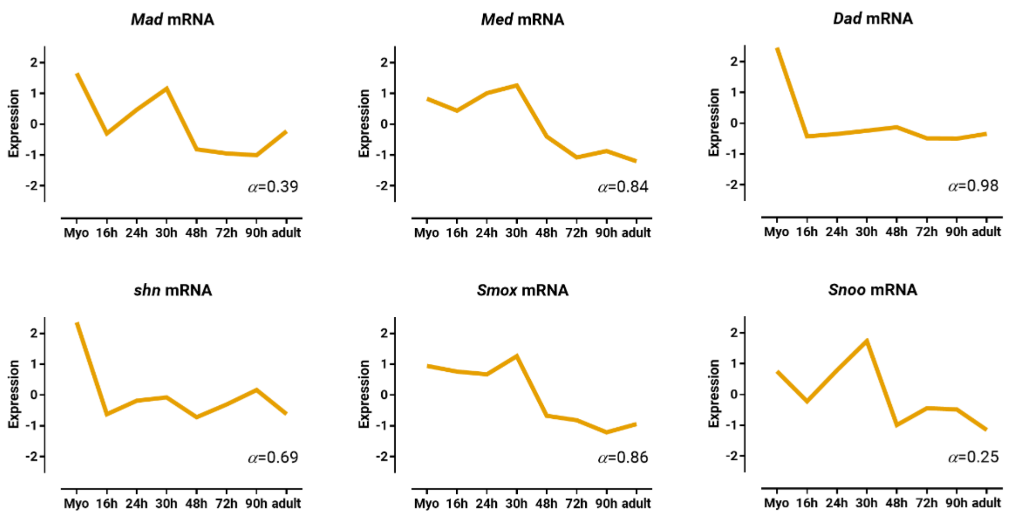


**Supplemental Figure 2.** Area proportional Venn diagrams representing transcriptional regulators binding to the 1 kb-long **(A)** and 5 kb-long **(D)** promoter regions of *Rbfox1-long* (fuchsia), *Rbfox1-short* (aqua), and *mir-33* (yellow). **(B)** List of transcriptional regulators common to the 1 kb-long promoter regions of *Rbfox1-long*, *Rbfox1-short*, and *mir-33*. Standard normal count values for **(C)** *Mad*, *Med*, *shn*, *Smox*, **(E)** *Dad*, and *Snoo* from an mRNA-seq developmental time-course of wild-type IFMs (Spletter et al., 2018).

Mukherjee_Supplemental_Fig_S3

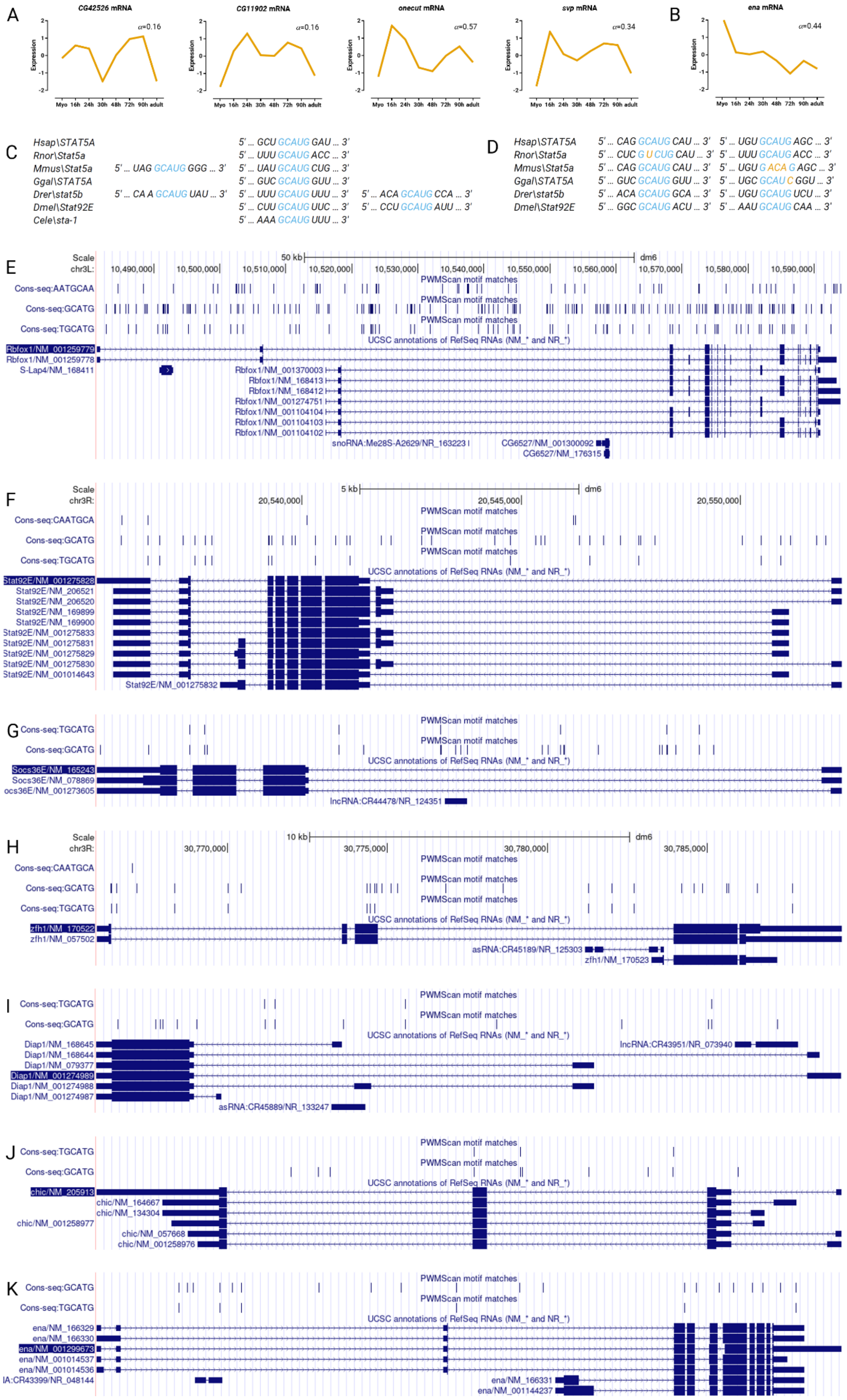

**Supplemental Figure 3.** Standard normal count values for **(A)** *CG42526*, *CG11902*, *onecut*, *svp*, and **(B)** *ena* from an mRNA-seq developmental time-course of wild-type IFMs (Spletter et al., 2018). Conserved binding sites of Rbfox1 at the **(C)** 3' end, and **(D)** 5' end *STAT* transcripts. Views near the **(E)** *Rbfox1*, **(F)** *Stat92E*, **(G)** *Socs36E*, **(H)** *zfh1*, **(I)** *Diap1*, **(J)** *chic*, and **(B)** *ena* loci captured from UCSC Genome Browser assembly ID: dm6. The location of Rbfox1 (*TGCATG* or *GCATG*), and *mir-33* (*AATGCAA* or *CAATGCA*) target sequences are noted.

A Stat92E-PK (UniProt: A0A0B4KH10)

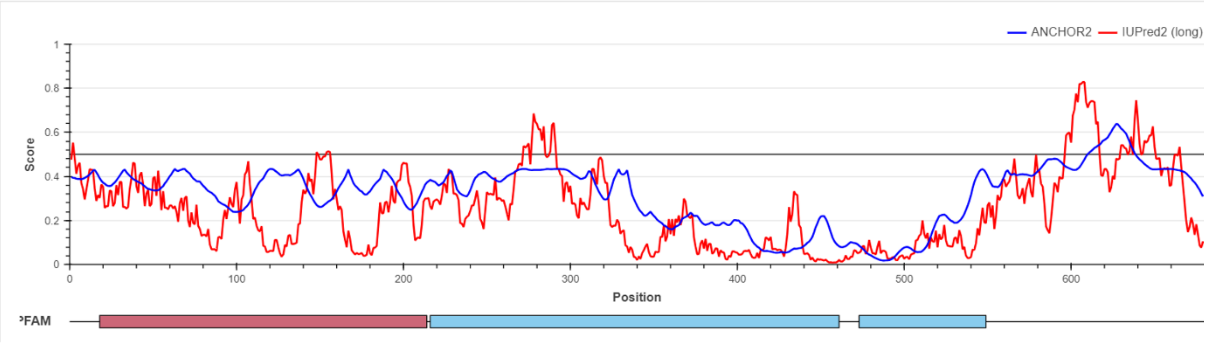


B Stat92E-PL (UniProt: A0A0B4KGE1)

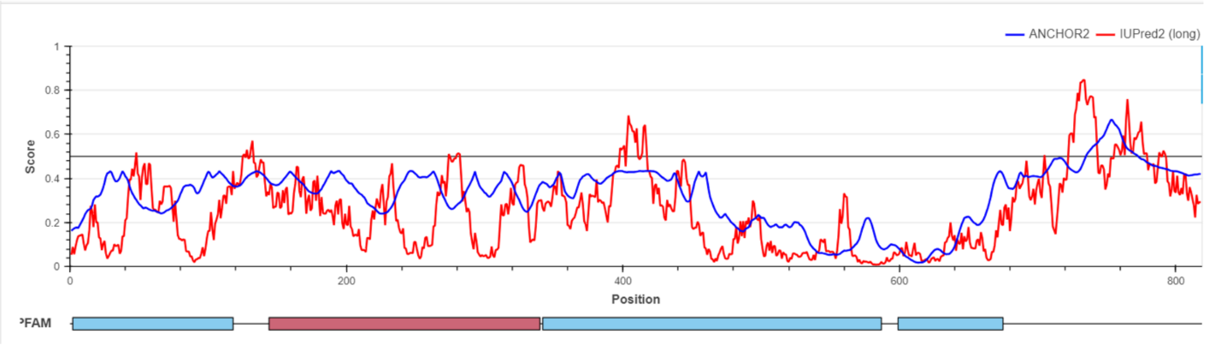


C Stat92E-PK (UniProt: A0A0B4KH10)

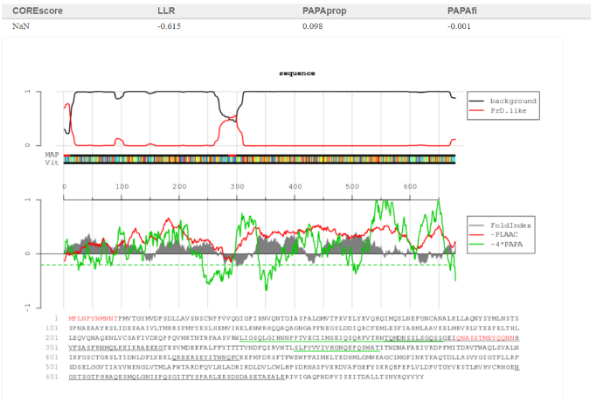


D Stat92E-PL (UniProt: A0A0B4KGE1)

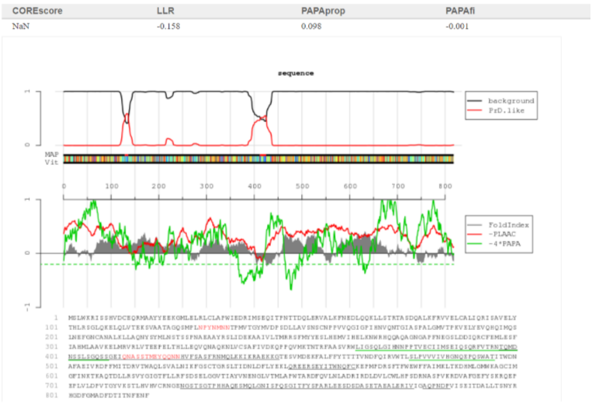


**Supplemental Figure 4.** Context-dependent predictions (default ANCHOR2) of IUPred2 long disorder (default) for **(A)** Stat92E-PK and **(B)** Stat92E-PL sequences. Identification of probable prion subsequences using PLAAC in **(C)** Stat92E-PK and **(D)** Stat92E-PL sequences, with Core Length=60, and Relative weighting of background probabilities ($\alpha$)=0 (meaning all from *Drosophila melanogaster*).

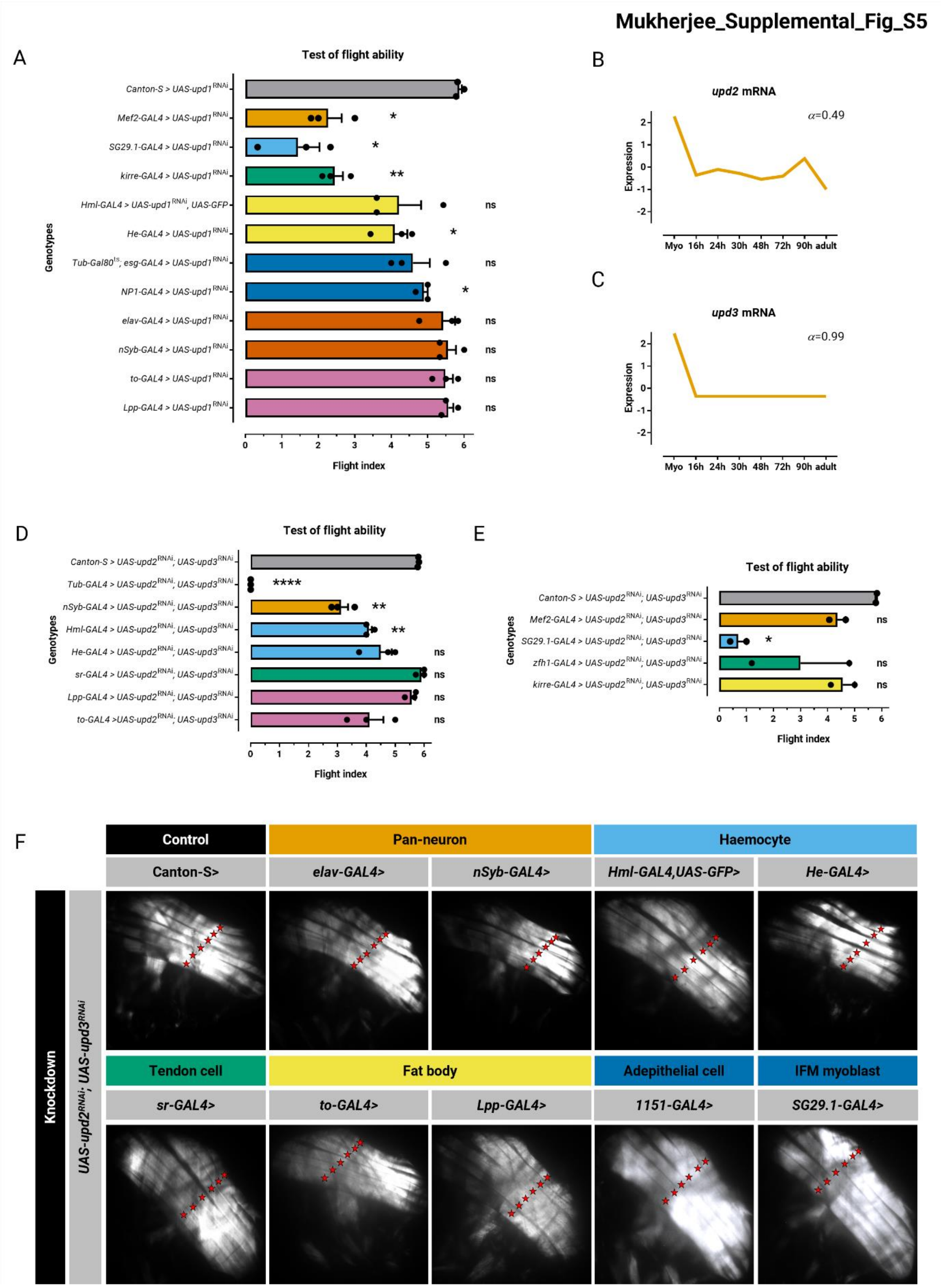


**Supplemental Figure 5.** Quantification of flight ability after **(A)** *upd1*, and **(D, E)** *upd2;upd3* knock-down. Genotypes as noted. Significance is from paired *t* test (**P* < 0.05; ***P* < 0.01). Standard normal count values for **(B)** *upd2*, and **(C)** *upd3* from an mRNA-seq developmental time-course of wild-type IFMs (Spletter et al., 2018). *Rbfox1* and *Stat92E* have similar temporal expression profiles. **(F)** Polarized microscopy images of hemithorax from flies. Genotypes as noted. Red stars indicate DLM fascicles.

**Mukherjee_Supplemental_Fig_S6**

A Zfh1-PB (UniProt: P28166-1)

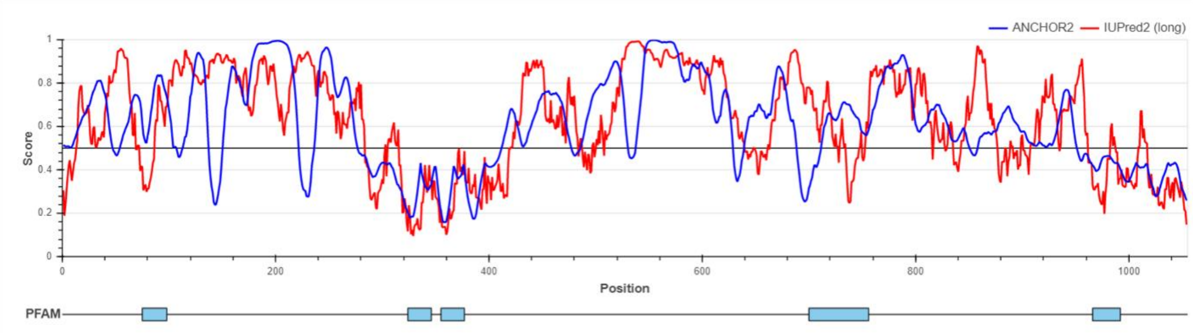


B Zfh1-PA (UniProt: P28166-2)

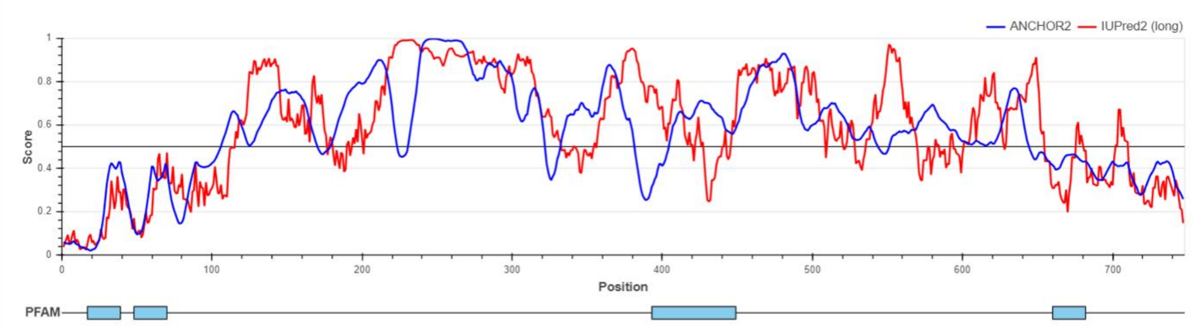


C Zfh1-PB (UniProt: P28166-1)

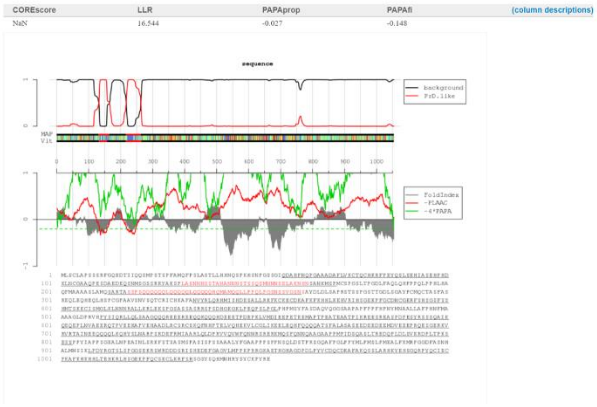


D Zfh1-PA (UniProt: P28166-2)

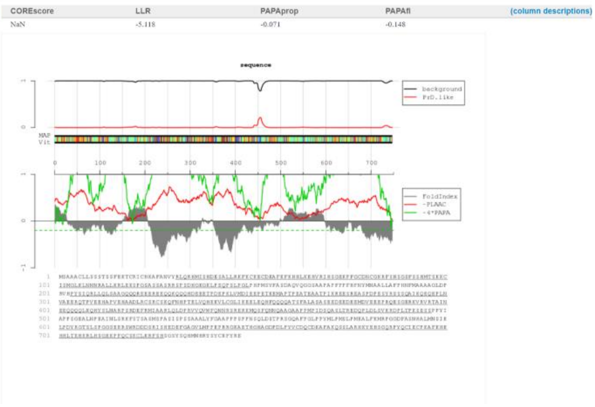


**Supplemental Figure 6.** Context-dependent predictions (default ANCHOR2) of IUPred2 long disorder (default) for **(A)** Zfh1-PB and **(B)** Zfh1-PA sequences. Identification of probable prion subsequences using PLAAC in **(C)** Zfh1-PB and **(D)** Zfh1-PA sequences, with Core Length=60, and Relative weighting of background probabilities ($\alpha$)=0 (meaning all from *Drosophila melanogaster*).

Mukherjee_Supplemental_Fig_S7

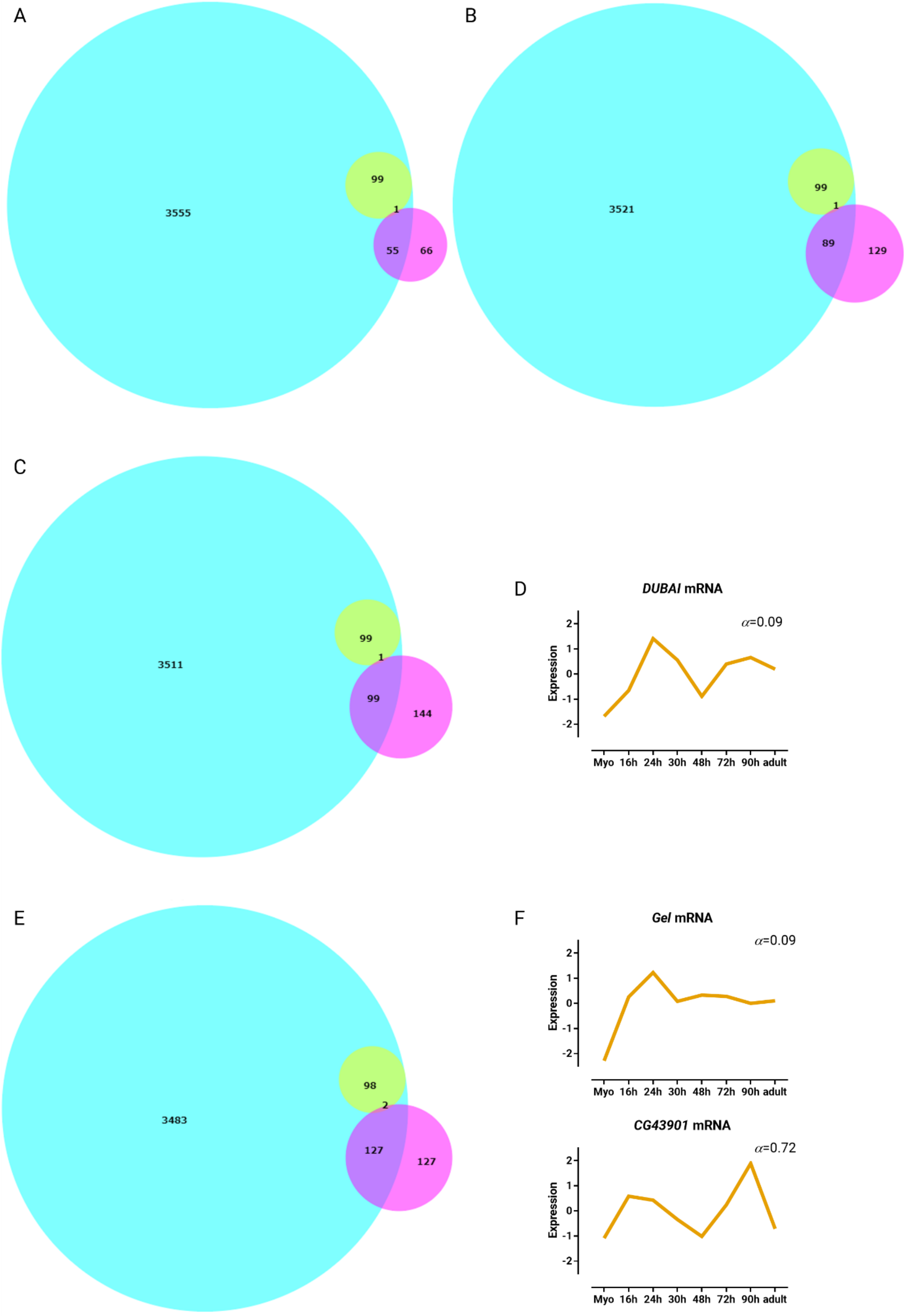

**Supplemental Figure 7.** Area proportional Venn diagrams representing all mapped putative *mir-33* (fuchsia) and Rbfox1 (aqua) targets, and Rbfox1 targets in Cluster 36 (yellow) relating to the GO biological process terms **(A)** ‘cell population proliferation’, **(B)** ‘regulation of cell population proliferation’, **(C)** ‘regulation of programmed cell death’, and **(E)** ‘actin filament organization’, respectively. Standard normal count values for **(D)** *DUBAI*, and **(F)** *Gel*, and *CG43901* from an mRNA-seq developmental time-course of wild-type IFMs (Spletter et al., 2018).
